# A defined platform of human peri-gastrulation-like biological fate patterning reveals coordination between Reaction-Diffusion and Positional-Information

**DOI:** 10.1101/102376

**Authors:** Mukul Tewary, Joel □stblom, Nika Shakiba, Peter W. Zandstra

## Abstract

How position dependent cell fate acquisition occurs during embryogenesis has been a central question in developmental biology. To study this process, we developed a defined, high-throughput assay using BMP4 to induce peri-gastrulation-like fate patterning in geometrically constrained human pluripotent stem cell colonies. We observed that, upon BMP4 treatment, phosphorylated SMAD1 (pSMAD1) activity in the colonies organized into a radial gradient – an observation mechanistically compliant with a BMP4-NOGGIN Reaction-Diffusion (RD) model. Consequent fate acquisition occurred as a function of both the pSMAD1 signaling strength, and induction time – consistent with the Positional-Information (PI) paradigm. Our findings implicate coordination between RD and PI underlying the peri-gastrulation-like fate patterning. This model not only predicts experimental results of perturbing key parameters like colony size, and BMP4 dose, but also identifies experimental conditions that rescue patterning in colonies of sizes that have been reported to be patterning-reticent, and recapitulate RD-like periodic patterns in large colonies.

## Introduction

During development, pluripotent stem cells (PSCs) in the epiblast are exposed to signaling gradients that initiate a sequence of fate specifications and cell movements resulting in the formation of spatially segregated germ layers in a developmentally conserved process called gastrulation^1–5^. A number of model systems have been used to study the molecular mechanisms that underpin the cell fate choices and morphogenetic events involved in the spatial ordering that arises during this critical developmental checkpoint^6–12^. However, it has been challenging to relate the regulatory mechanisms that generate pattern formation in other developmental model systems to human gastrulation.

Studying human gastrulation-associated pattern formation requires a platform for the robust simulation and investigation of the signaling programs that initiate and drive gastrulation-like events. We and others have previously used micro-patterning technologies to control human (h)PSC colony geometry, demonstrating increased cell response consistency^13–19^. These studies highlight that micro-environmental control of endogenous signaling profiles, cell-cell contact, and mechanical forces are crucial to robustly regulate cell fate and spatial tissue organization. Warmflash *et al*.^20^ used similar techniques to demonstrate that following BMP4 treatment, geometrically-controlled hPSC colonies could recapitulate many aspects of a peri-gastrulation-stage epiblast. These colonies exhibited spatially patterned regions characteristic of primitive-streak-like, mesodermal, endodermal, ectodermal and trophoblast-like tissues^20^. Recently, Etoc *et al.*^21^ suggested that this spatial patterning is mediated by the self-organized formation of a BMP signaling gradient. Mechanistically, they implicated a combination of BMP signaling induced expression of NOGGIN in hPSCs, and decreased sensitivity to BMP4 due to reduced accessibility to BMP receptors at the center of the colonies, and speculated that the fate patterning mediated by this signaling gradient could rely on a threshold model.

Two prominent biochemical models have influenced our understanding of cell fate patterning and morphogenesis that occurs during embryonic development: reaction-diffusion (RD) – proposed by Alan Turing^22–24^; and positional information (PI) – proposed by Lewis Wolpert^22,25,26^. The RD model predicts the presence of an interaction network between two diffusible interacting molecules – an ‘activator’, which activates the expression of both molecules, and an ‘inhibitor’, which inhibits their expression. Turing proposed that initiation of this interaction network, together with dissimilar diffusivities, would be sufficient to give rise to complex biological pattern formation. In contrast, the PI model, asserted that biological spatial patterns require prior asymmetries in morphogen distribution across a developing tissue; and that the asymmetries are of a scale that would allow cells to reliably specify fates through a signaling threshold-based mechanism. Studies using the developing neural tube as an experimental system have demonstrated that the PI paradigm was not just mediated by morphogen signaling threshold levels; a relationship between the morphogen concentration the length of time of morphogen exposure also influences cells fates ^22,27–29^.

Here we demonstrate, using fully defined and scalable (96-well plate) conditions, that geometrically-specified hPSC colonies organize into radially segregated regions that express markers characteristic of ectoderm-like, primitive streak-like, and trophoblast-like tissues. We show that upon BMP4 induction, a BMP4-NOGGIN RD network rapidly organizes pSMAD1 activity into a signaling gradient within the geometrically confined colonies. The established gradient then patterns PSC differentiation in a manner consistent with the PI paradigm. Further, we demonstrate that, across a range of colony sizes and BMP4 doses, the RD model consistently predicts the formation of the pSMAD1 signaling gradient, and the PI paradigm accurately predicts the patterned fate acquisition. We also identify previously unreported periodic patterns of fate patterning consistent with the RD paradigm, and rescue fate patterning in colonies that have been thought of as being incapable of facilitating pattern formation. Taken together, our data support the concept that a two-step process, involving coordination between the RD and PI models, directs the peri-gastrulation-like patterns that develop in differentiating hPSC colonies.

## Results

### A defined high throughput assay for induction of peri-gastrulation-like events in human pluripotent stem cell colonies

It has previously been reported, and reproduced in our hands (**Fig 1a**), that radially segregated expression of peri-gastrulation-associated markers CDX2, Brachyury (BRA), and SOX2 - representative of trophoblast-like, primitive-streak-like, and ectoderm-like tissues respectively – develop in geometrically-controlled, 1000 µm diameter circular hPSC colonies differentiated in BMP4 supplemented mouse embryonic fibroblast conditioned medium (CM)^20,21^. Given that identification of key molecular regulators of the peri-gastrulation-like pattern formation in the presence of undefined CM is difficult, our first aim was to identify defined basal conditions that induce these patterning events in hPSC colonies. Further, we optimized a previously described protocol that uses Deep Ultraviolet light (< 200 nm) mediated photo-oxidation of polyethylene glycol (PEG) coated slides^30^ to allow high-fidelity patterning of hPSC colonies and adapted this technique to produce 96-well microtiter plates (Materials and Methods).

**Figure 1:**
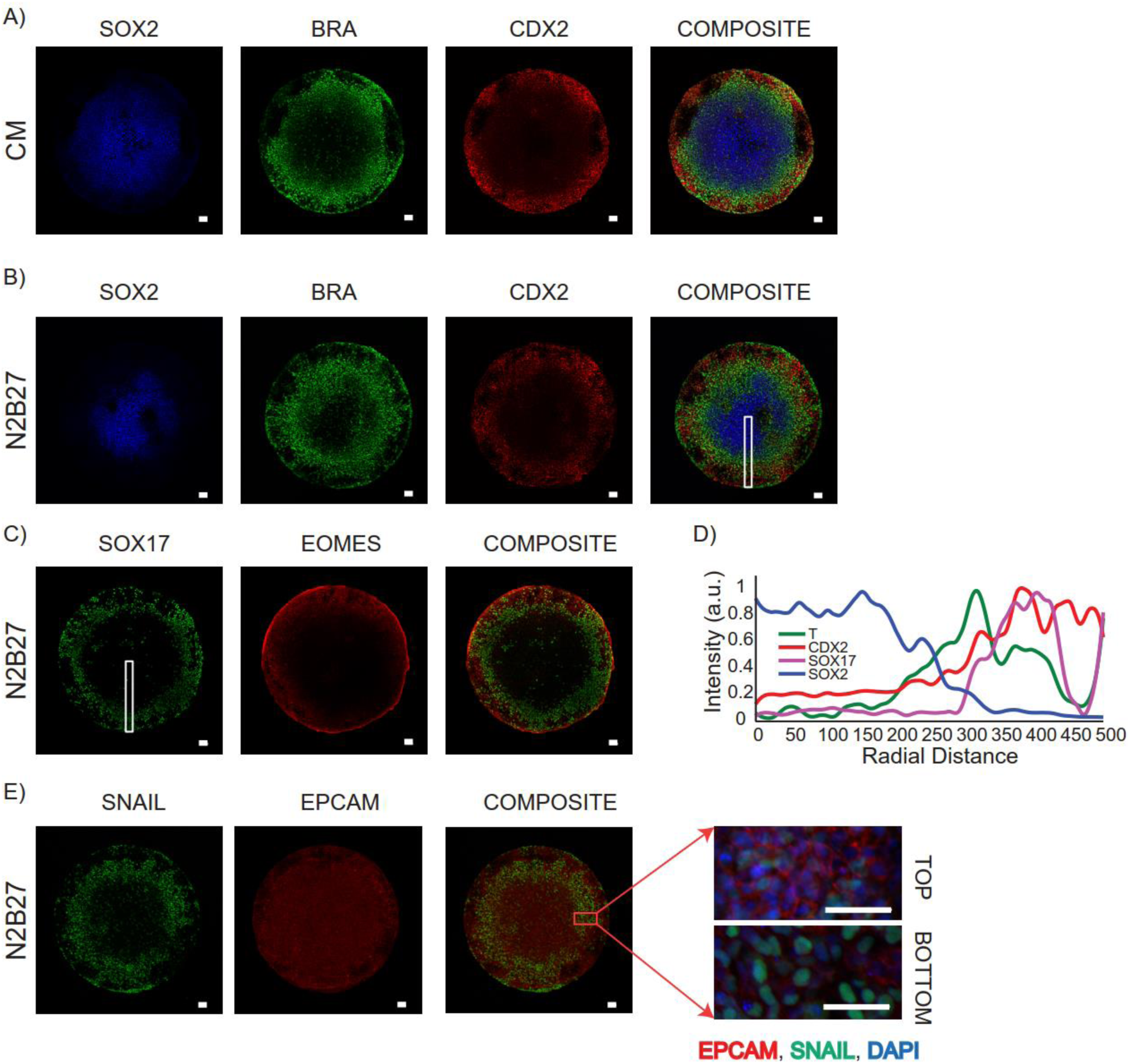
Defined peri-gastrulation-like patterning induction in differentiating hPSC colonies. Representative immunofluorescence images of: A) fate patterning of SOX2, BRA, and CDX2 in BMP4 supplemented CM as previously reported^20,21^; B) fate patterning in BMP4 supplemented N2B27 medium stained for SOX2, BRA, and CDX2, and C) SOX17, and EOMES. D) Spatial trends for intensity of expression of SOX2, BRA, CDX2, and SOX17 in regions marked by white rectangle in B and C (Average trends of replicates shown in **Sup Fig 1, 2**). E) Representative image of SNAIL and EPCAM staining in micro-patterned colony differentiated in N2B27 shows that mesenchymal marker expressing cells in the primitive streak region are located underneath an epithelial layer. Scale bars represent 50 µm.

Using our PEG plates, we set out to screen the following media: Nutristem (NS), mTeSR (MT), Essential-8 (E8), a Knockout Serum Replacement based medium (SR), and an N2B27 based medium (N2B27), supplemented with BMP4, for induction of peri-gastrulation-like patterning in geometrically defined hPSC colonies. Preliminary testing with N2B27 media revealed a positive response in the induction of the BRA expressing region to NODAL supplementation (data not shown), which is consistent with the importance of Nodal signaling in the induction of the primitive streak^31^. Therefore, we performed all following N2B27 based experiments with the media supplemented with 100ng/ml of NODAL. We found that, although segregated expression of CDX2, and SOX2 developed in all defined test media, BRA expression was not consistently observed in E8, NS, and SR media (**Sup Fig 1, 2**). Further, while reproducible induction of peri-gastrulation-like patterns developed in both N2B27 and MT media, colonies patterned in MT tended to lift off the plate by 48h. BMP4 supplemented N2B27 medium robustly gave rise to the differentiating hPSC colonies with regions expressing trophoblast-associated (CDX2) and primitive-streak-associated (BRA) markers, with concurrent spatial patterning of the germ layers – endoderm (SOX17), mesoderm (EOMES) and ectoderm (SOX2), in a manner indistinguishable from hPSC colonies differentiated in BMP4-supplemented CM (**Fig 1b-d, Sup Fig 1, 2**)^20^. Notably, the SNAIL-expressing mesenchymal cells appeared underneath the EPCAM-expressing epithelial layer in the primitive streak marker-expressing regions of our colonies, (**Fig 1e**), consistent with observations reported in BMP4-supplemented CM^20^. Given these results, we selected N2B27 medium for our platform to study *in vitro* peri-gastrulation-like patterning in hPSC colonies.

### BMP4-NOGGIN interaction network regulates pSMAD1 gradient self-organization

As our N2B27 media was supplemented with NODAL to ensure reproducible induction of the primitive-streak-like region, and Nodal signaling is involved in the induction of the primitive streak in differentiating hPSCs^20,31^, we first asked if NODAL was involved in the initiation of the morphogenetic events as well. We found that although Nodal signaling was indispensable for the induction of the BRA expressing region (**Sup Fig 3a**), in the absence of BMP signaling, NODAL was unable to induce the peri-gastrulation-like events (**Sup Fig 3a, b**). On the other hand, BMP signaling could initiate the morphogenetic reorganization even in the absence of Nodal signaling (**Sup Fig 3c-e**). Consequently, we focused on investigating how BMP signaling induced the morphogenetic changes and peri-gastrulation-like events in the differentiating hPSC colonies. As a first step, we focused on identifying how the downstream effectors of BMP signaling were organized within the micro-patterned colonies. BMP ligands mediated activation of the BMP receptors (BMPR1A, BMPR1B, BMPR2) results in phosphorylation and nuclear translocation of SMAD1 followed by the transcription of context-specific target genes^32^. We measured the dynamics of pSMAD1 signaling as a function of the radial distance from the center of the differentiating colonies at 1, 6, 18, and 24 h following BMP4 induction. Robust analysis was enabled by overlaying the pSMAD1 expression profiles of at least 100 colonies at each time point, yielding the average intensity of pSMAD1 as a function of the colony radius. We noted that an hour after BMP4 induction, the pSMAD1 activity appeared evenly at all colony radii (**Fig 2a**). This implies that signaling capabilities of the cells immediately after the BMP4 presentation are position-independent. Subsequently, over the first 24 h of BMP4 treatment, pSMAD1 activity appeared to spontaneously organize into a radial gradient with the cells in the periphery experiencing higher levels of pSMAD1 activity (**Figs 2a-c, and Sup Fig 4**). To test whether the self-organization of pSMAD1 activity could be attributed to regulatory feedback of the BMP pathway, we measured the expression of positive and negative feedback mediators of BMP signaling at over the first 26 h after BMP4 treatment. We found significant upregulation in the expression of both NOGGIN, and BMP4 (**Fig 2d**) in a dose dependent manner (**Sup Fig 5b**). Notably, BMP4 and NOGGIN upregulation in response to BMP4 treatment was seen in all basal medium conditions tested (**Sup Fig 6**), indicating that the upregulation was a response to BMP signaling and not related to the N2B27 medium. To specifically test whether NOGGIN is involved in the self-organization of the pSMAD1 gradient, we evaluated the effect of NOGGIN inhibition on the pSMAD1 gradient formation. NOGGIN inhibition, using SiRNA, significantly increased pSMAD1 activity in the central regions of the micro-patterned colonies demonstrating a role for NOGGIN in the self-organization of the pSMAD1 gradient (**Fig 2e-g, Sup Fig 7**), as has recently been proposed by others^21^. Interestingly, along with the NOGGIN mediated negative feedback, we noted the presence of a positive feedback loop in BMP signaling wherein BMP4 supplementation resulted in BMP4 expression (**Fig 2d, Sup Fig 6**). Together, the presence of both a positive and a negative feedback mediated by BMP4 implicate a BMP4-NOGGIN RD network orchestrating the pSMAD1 radial self-organization in differentiating hPSC colonies (**Fig 2h**). We next set out to investigate properties of the RD framework to determine how patterning of hPSC-derived peri-gastrulation-like events arise.

**Figure 2:**
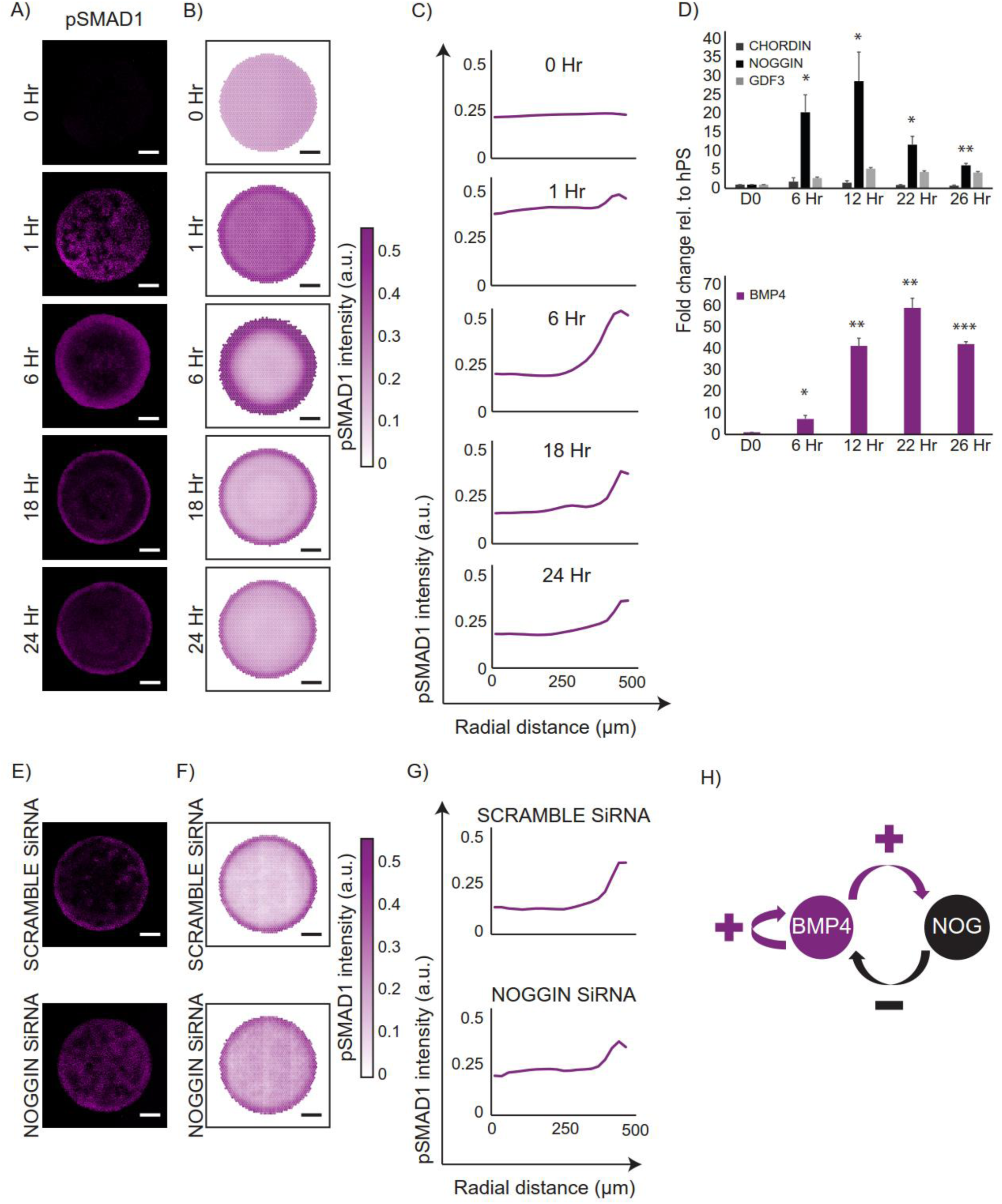
pSMAD1 gradient self-organization in differentiating colonies implicates a BMP4-NOGGIN RD network. A) Representative immunofluorescence images of colonies stained for pSMAD1 at 0, 1, 6, 18, and 24 hours after BMP induction. B) Average radial gradient of pSMAD1 signaling represented as overlays for 202, 193, 105, 100, 105 colonies for the respective induction times. Data collected from two experiments. C) The average radial trends of pSMAD1 activity at each induction duration. D) Temporal gene expression profiles for BMP signaling inhibitors (CHORDIN, NOGGIN, and GDF3) and BMP4 at 6, 12, 22, and 26 hours. Data shown as mean and standard deviation (S.D.) of three independent experiments. * p<0.05, ** p<0.01, *** p<0.001. E) Representative pSMAD1 immunofluorescence images for colonies treated with scramble SiRNA and NOGGIN SiRNA. F) Overlays of colonies treated with Scramble siRNA, NOGGIN SiRNA (64, and 62 colonies respectively). Data collected from two experiments. G) Average radial trends of pSMAD1 signaling dynamics for the Scramble, and NOGGIN SiRNA conditions. H) Overview of the implicated BMP4-NOGGIN RD network. Scale bars represent 200 µm.

### pSMAD1 gradient formation is colony size and BMP4 concentration dependent

To simulate RD mediated self-organization of pSMAD1 activation pattern, we developed a finite element model that predicts the spatial and temporal distribution of signaling-competent, free BMP4 ligands within the differentiating geometrically confined colonies using the RD-specific two-component, coupled, partial differentiation equation set^23,33^ (**Supplementary Information, Sup Video 1**). To determine if this model could accurately predict experimental data, we performed sweeps on two key parameters that regulate the formation of the free BMP4 distribution within the hPSC colony: the initial concentration of BMP4 in the induction media (BMPi), and the colony size. Our model predicted that reducing BMPi while maintaining the colony diameter at 1000µm would still lead to the formation of free BMP4 distribution gradients within the colonies, but with lower ligand levels at the colony periphery (**Fig 3a-b**). Consistent with model predictions, when we tested the spatial dynamics of the pSMAD1 concentration gradients formed in 1000µm diameter colonies 24 h after induction with varying BMPi (6.25 ng/ml, 12.5 ng/ml, 25 ng/ml, and 50 ng/ml), we observed the formation of pSMAD1 activity gradients at all concentrations, with lower BMPi conditions producing lower levels of peak pSMAD1 activity at the radial edge of the colonies (**Fig 3c-e**, Sup Fig 8). We next queried the model to predict how reducing colony size, at a fixed BMPi dose, would affect the spatial dynamics of free BMP4 ligands in the differentiating colonies. The model predicted that reducing the colony size would progressively increase the presence of free BMP4 ligands at the center of the colonies (**Fig 3f-g**). To test the model predictions experimentally, we differentiated colonies of varying sizes with constant BMPi dose (50ng/ml), and assessed pSMAD1 activity 24 h post-induction. We found that small colonies were unable to form regions with low pSMAD1 signaling (**Fig 3h-j, Sup Fig 9**). These findings are consistent with our hypothesis that the self-organization of the pSMAD1 pattern activity observed in differentiating hPSC colonies is governed by a BMP4-NOGGIN RD network.

**Figure 3:**
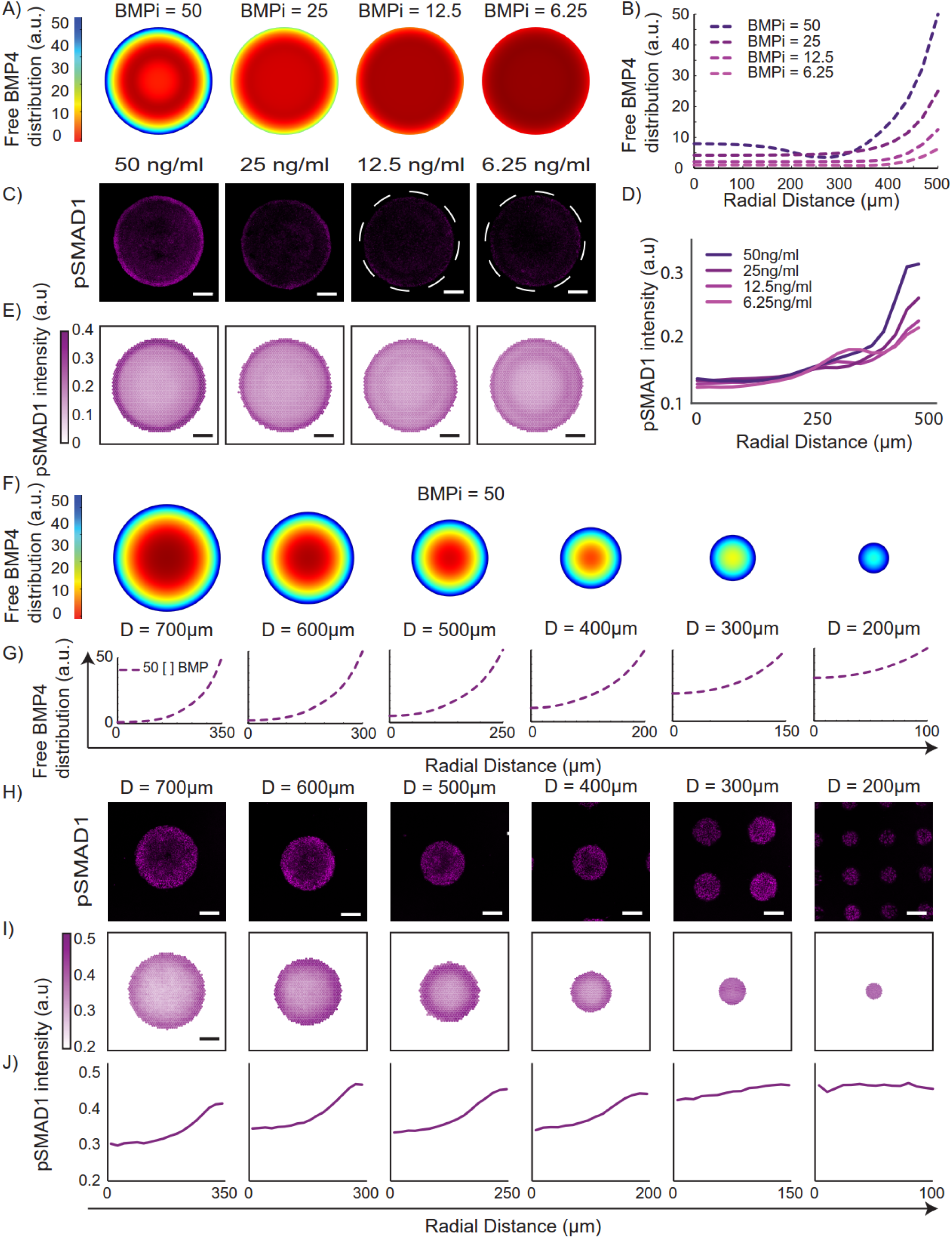
BMP4-NOGGIN RD model predicts pSMAD1 gradient response to colony size and BMP4 dose perturbations. A) Predicted distribution of free BMP4 ligands in colonies of 1000 µm diameter as a function of varying BMPi concentration (50 ng/ml, 25 ng/ml, 12.5 ng/ml, and 6.25 ng/ml). B) Graphical depiction of predicted distributions. C) Representative immunofluorescence images of micro-patterned colonies stained for pSMAD1 24 hours after induction with varying BMPi conditions (50 ng/ml, 25 ng/ml, 12.5 ng/ml, and 6.25 ng/ml). D) pSMAD1 signaling dynamics represented as a function of the colony radius and E) overlays of 143,119, 129, 163 colonies for respective conditions. The results were collected from two experiments. F) Predicted distribution of free BMP4 ligands, following induction with 50 ng/ml BMP4, as a function of colony diameter (700 µm, 600 µm, 500 µm, 400 µm, 300 µm, and 200 µm). Graphical depiction of predicted distributions in F. H) Representative immunofluorescence images of colonies stained for pSMAD1 24 hours after induction with 50 ng/ml of BMP4. I) Overlays of 87, 118, 178, 261, 437, and 373 colonies for respective conditions. J) pSMAD1 signaling dynamics represented as a function of the colony radius. The results were collected from two experiments. Scale bars represent 200 µm.

### BMP signaling gradient patterns fate via PI paradigm

Notably, the perturbations we tested above resulted in a change in pSMAD1 activity both at the periphery (**Fig 3d**) and at the center (**Fig 3j**) of the colonies. We reasoned that if the fate acquisition in these colonies was a function of pSMAD1 activity thresholds, we would observe fate switches at different conditions. Accordingly, we tested the same conditions above – this time after 48 h – and stained for the fate-associated markers: SOX2, BRA, and CDX2. First, to perturb the pSMAD1 activity at the colony periphery, we varied BMPi doses while keeping the colony size constant (**Fig 4a**). Consistent with a threshold-dependent fate acquisition model, we found a significant reduction of CDX2 expression in colonies induced at low BMPi conditions (**Fig 4b-c**). Furthermore, when we perturbed the pSMAD1 activity at the colony center by varying colony size at a constant BMPi dose (**Fig 4d**), we found that SOX2 expression was absent in smaller colonies (< 400 µm diameter) (**Fig 4e-f**). These data indicate that the fate acquisition in the differentiating geometrically-confined hPSC colonies is suggestive of BMP signaling thresholds.

**Figure 4:**
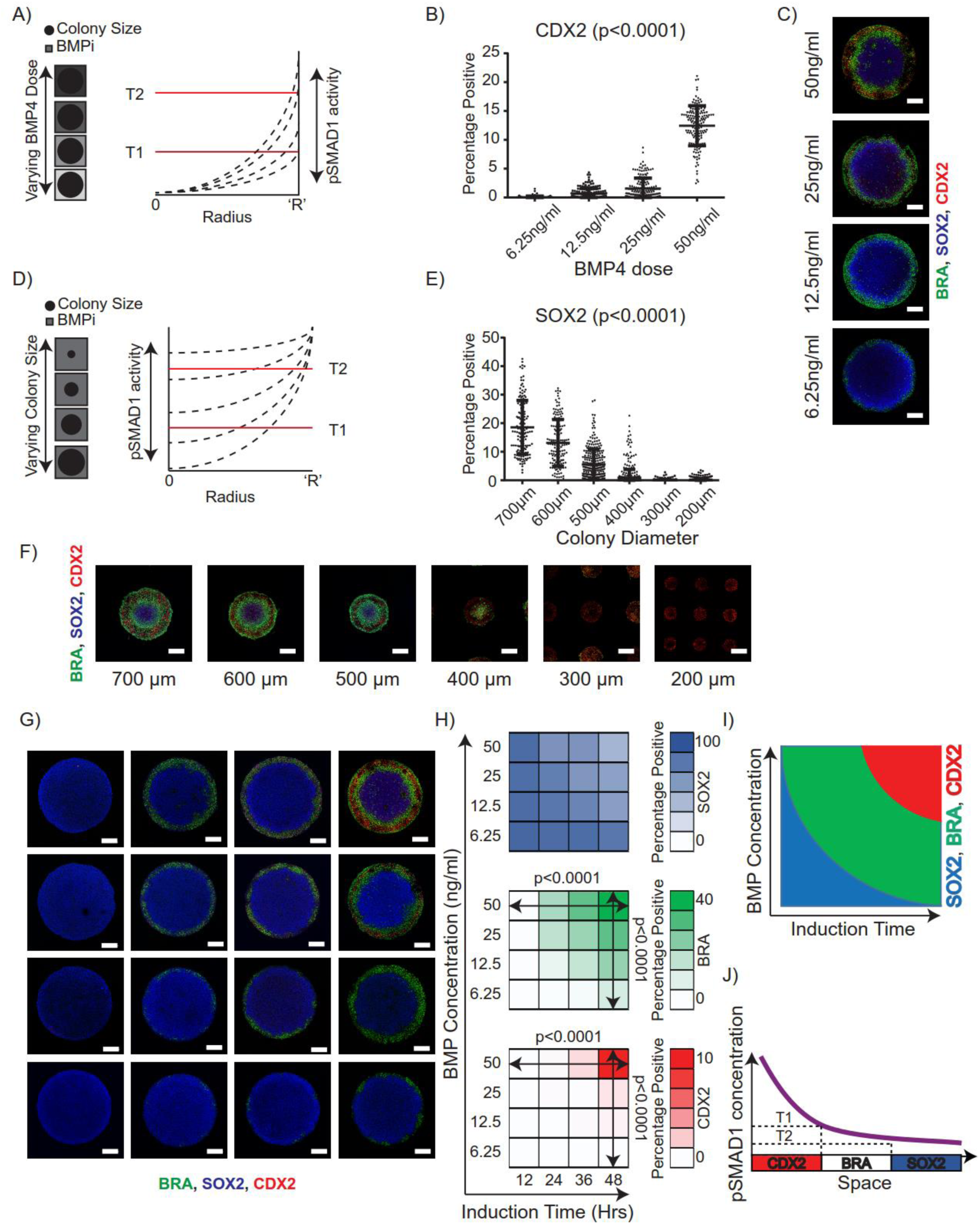
Fate patterning in hPSC colonies arises in a pSMAD1 threshold dependent manner. A) Overview of the experimental setup: BMPi is varied while maintaining colony size constant to perturb pSMAD1 signaling at the colony periphery. B) Quantification of cells expressing CDX2 in 1000 µm colonies induced to differentiate at varying BMPi concentrations (6.25ng/ml, 12.5ng/ml, 25ng/ml, and 50ng/ml) (p value calculated using one way ANOVA). Number of colonies are 136, 168, 169, and 156 for the respective conditions. Results pooled from two separate experiments. Data represented as mean (s.d). C) Representative immunofluorescence images of SOX2, BRA, and CDX2 expression in geometrically confined 1000 µm diameter colonies differentiated in 6.25ng/ml, 12.5ng/ml, 25ng/ml and 50ng/ml of BMP4. D) Overview of the experimental setup: colony size is varied while maintaining constant BMPi perturbs the level of pSMAD1 signaling in the colony center. E) Quantification of cells expressing SOX2 in colonies of varying diameters (700 µm, 600 µm, 500 µm, 400 µm, 300 µm, 200 µm) differentiated in BMPi = 50ng/ml (p value calculated using one way ANOVA). Number of colonies analyzed from two separate experiments were 144, 160, 279, 466, 789, and 1607 for the respective conditions. Data represented as mean (s.d). F) Representative immunofluorescent images of SOX2, BRA, and CDX2 expression in geometrically confined colonies of varying diameters (700µm, 600µm, 500µm, 400µm, 300µm, and 200µm). G) Representative immunofluorescence images of SOX2, BRA, and CDX2 stained 1000µm diameter colonies differentiated at varying BMPi concentrations (50ng/ml, 25ng/ml, 12.5ng/ml, and 6.25ng/ml) and induction times (12, 24, 36, and 48 hours). H) Average percentage of cells expressing SOX2, BRA, and CDX2 in the colonies. Each condition had over 140 colonies. Data pooled from two experiments. The p values were calculated using one way ANOVA. I) Overview of peri-gastrulation-like fate acquisition arising as a function of both morphogen concentration and induction time. J) Overview of the fate acquisition arising in a manner consistent with Positional Information. Scale bars represent 200µm.

Signaling threshold-mediated fate acquisition is reminiscent of the positional information (PI) model of cell biological fate patterning^25,26^ (i.e., different signaling thresholds induce transcription factor (TF) networks associated with different cell fates). However, in typical models that pattern fates via the PI paradigm the TFs associated with patterned fates arise not just as a function of the morphogen concentration, but also as a function induction time (i.e. the fates associated with higher levels of morphogen concentration are induced after longer induction times^22,27–29^). Therefore, we set out to investigate if the fates in the differentiating hPSC colonies arose as a function of both the level of pSMAD1 activity and the time of induction. Accordingly, we tested four different BMPi doses (50 ng/mL, 25 ng/mL, 12.5 ng/mL, and 6.25 ng/mL), and analyzed the patterned fates that emerged at four different induction times (12h, 24h, 36h, and 48h). We found that the BRA and CDX2 fates did not arise consistently at either lower BMPi doses or at shorter durations of higher BMPi doses (**Fig 4g-h, and Sup Fig 10**). Taken together, this analysis suggests that the development-relevant cell fate patterning in the hPSC colonies mediated by the self-organized pSMAD1 gradient follows the PI paradigm, as reflected by the characteristic PI-like profile (**Fig 4i**) produced in a plot of induction interval against BMP4 concentration. In summary, our findings demonstrate that the fate patterning within the differentiating hPSC colonies occurs via the cardinal positional information model (**Fig 4j**).

### A two-step process for peri-gastrulation-like biological pattern formation

Our data indicate that the formation of the pSMAD1 gradient pattern follows a RD mechanism, and the subsequent fate acquisition follows a cardinal PI mechanism; together resulting in a two-step process of biological fate patterning in the geometrically confined hPSC colonies. Recognizing that the classic RD models have repetitive and periodic peaks of fate signaling activity^22–24^, we set out to test the two-step model of biological fate patterning by querying our model to identify conditions that would result in a periodic specification of patterned fates. Our model predicted that although an increase in the colony size alone would be insufficient to induce a periodic response in presence of free BMP4 ligands, a concomitant increase in BMPi could capture that response (**Fig 5a-b, Sup Video 2-3**). Consistent with the prediction, pSMAD1 levels at 24h post induction appeared to have a periodic response in a 3mm diameter colony induced to differentiate in 200 ng/ml of BMP4, but not in 50 ng/ml of BMP4 (**Fig 5c-d, Sup Fig 11**). Furthermore, when the 3mm diameter colonies were stained for SOX2 and BRA at 48 h post induction, we observed a periodic response of BRA expression arise in the colonies differentiated in 200ng/ml of BMP4 (**Fig 5e-f, Sup Fig 12**).

**Figure 5:**
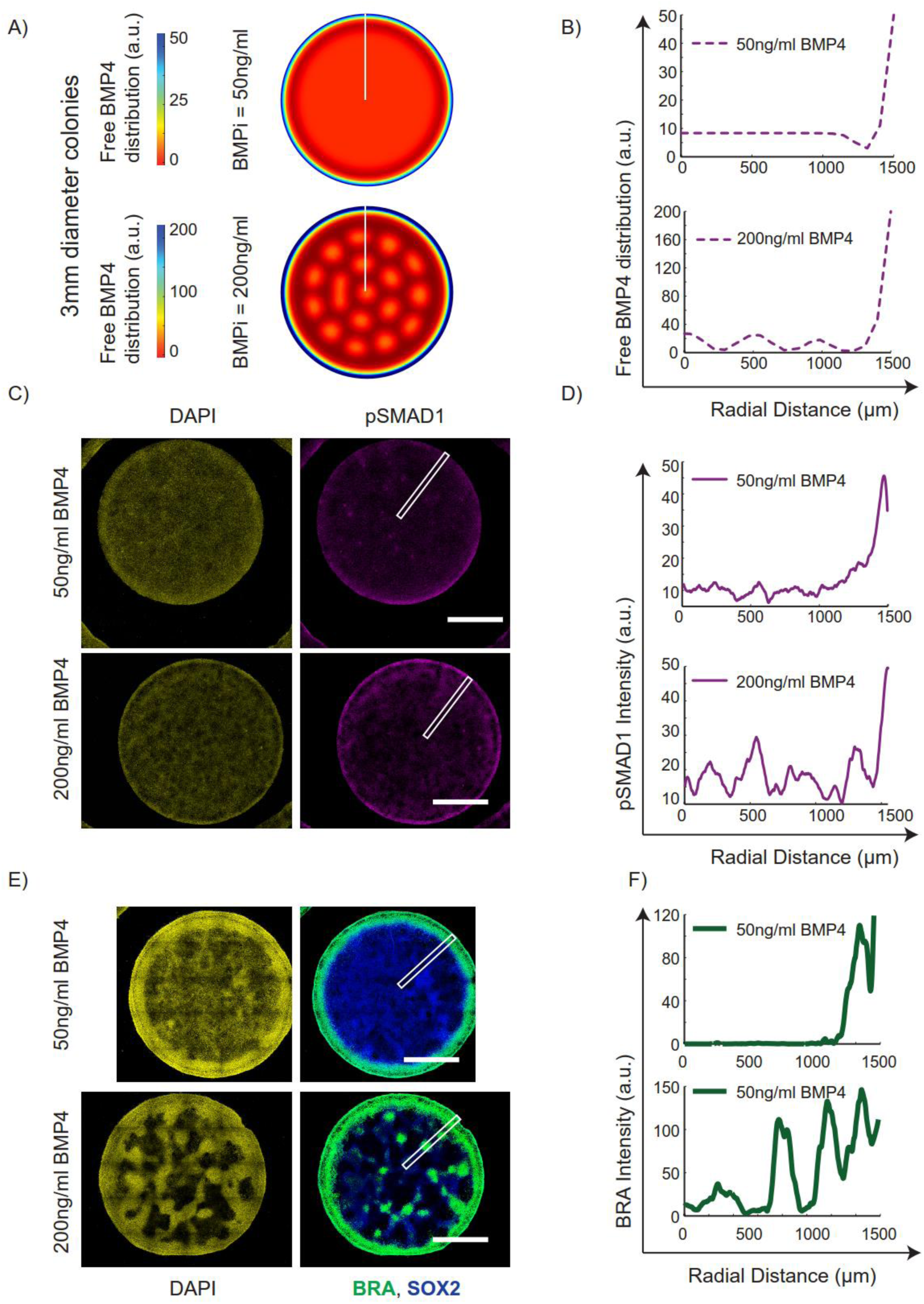
Fate patterning occurs via a unified mechanism of RD and PI in differentiating micro-patterned hPSC colonies. A) Model predictions for gradient formation of free BMP4 ligands indicate that higher BMPi concentrations would result in a typical RD-like periodic response of free BMP4 ligands. B) Line plot representation (measured along white lines in A) of free BMP4 ligands. C) Representative immunofluorescent images of nuclei and pSMAD1 staining at 24 hours of differentiation with 50ng/ml and 200ng/ml of BMP4. D) Line plots (along white rectangles in C) show RD-like response in pSMAD1 activity for BMPi = 200ng/ml. E) Representative immunofluorescent images of colonies differentiated for 48 hours with 50ng/ml and 200ng/ml of BMP4 stained for DAPI, BRA and SOX2. F) Line plot representation shows a cardinal RD-like behavior in the fate patterning of BRA for colony differentiated with 200ng/ml of BMP4. Scale bars represent 1mm.

As a final validation, we considered the claims of previous reports that have indicated a crucial need of edge sensing underlying the observed pattern formation^21^; a mechanism which renders small colonies of 250µm diameter reticent to fate patterning – only allowing the induction of the CDX2 expressing Trophoblast-like region^20^. Since our interpretation of the data mechanistically implicated a step-wise coordination of RD and PI, our interpretation of the inability of 250µm diameter colonies to induce all three fates was different from the above proposition. Specifically, we argue that at a BMPi dose of 50ng/ml, the RD mediated organization of free BMP4 ligands within colonies of 250µm diameter were sustained at high levels throughout the colony (**Fig 3F-J**), and past the thresholds that would induce the primitive-streak-like and the ectoderm-like region after a 48h induction (**Fig 4E-F**) as per the PI paradigm. To demonstrate this claim, we asked our model if perturbing the BMPi conditions could induce the organization of free BMP4 ligands at appropriate levels to rescue the fate patterning of all three lineages. Our model predicted that reducing BMPi would reduce the levels of free BMP4 ligands throughout the colony whereby the BRA, and SOX2 expressing regions could be rescued (**Fig 6A-B**). We tested the expression profiles of pSMAD1 activity at 24h post peri-gastrulation-like induction with varying BMPi conditions (50ng/ml, 25ng/ml, 12.5ng/ml, and 6.25ng/ml) of numerous colonies (over 500 per dose condition), and consistent with our model predictions, we observed a reduction of pSMAD1 levels overall – but especially in the center of the colonies (**Fig 6C-E, Sup Fig 13**). These data suggested that lower BMPi conditions might rescue the formation of the BRA, and SOX2 expressing regions in colonies of 250µm diameter in accordance with the PI paradigm. Indeed, when we tested the fate expression within 250µm diameter colonies after 48 hours of induction with varying BMPi doses, we observed a significant reduction of CDX2 expression with reduced doses of BMP4 with a concomitant increase of BRA and SOX2 expression within the colonies (**Fig 6F-G**). Importantly, at a BMPi dose of 12.5ng/ml, we see a rescue of patterning of all three fates within colonies of 250µm diameter (**Fig 6G**).

**Figure 6:**
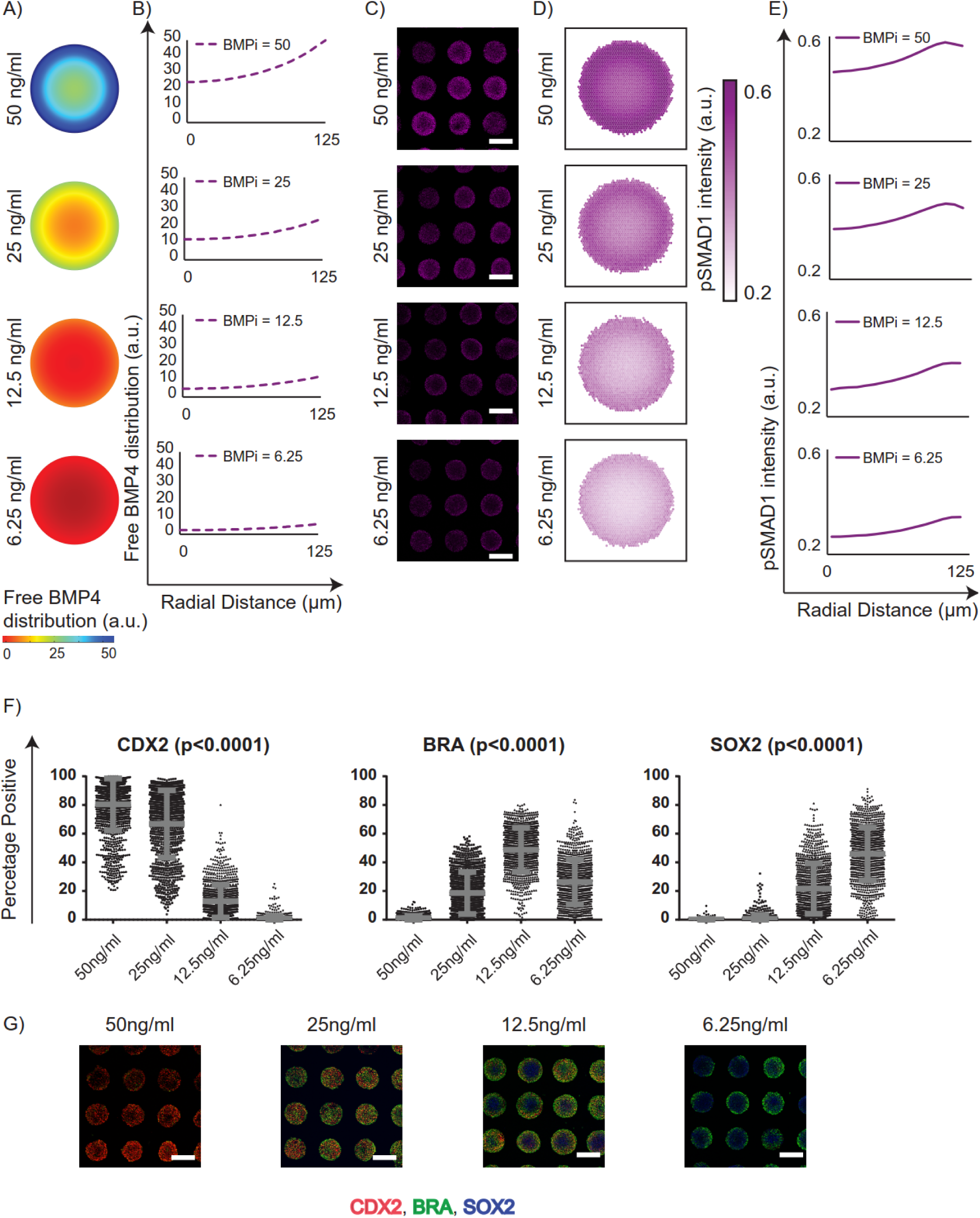
Reducing BMP4 dose in induction media rescues fate patterning in 250µm diameter colonies. A) Model predictions for gradient formation of free BMP4 ligands distribution in 250µm diameter colonies in response to varying BMPi doses. B) Line plot representation of free BMP4 ligands as a function of the colony radius. C) Representative immunofluorescence images of colonies stained for pSMAD1 24 hours after induction with varying doses of BMP4. D) Overlays of 976, 638, 475, and 689 colonies for respective conditions. E) Signaling dynamics represented as a function of the colony radius. The results were collected from two experiments. F) Quantification of percentage of cells in each colony expressing CDX2, BRA, and SOX2 in 250µm diameter colonies induced to differentiate at varying BMPi concentrations (6.25 ng/ml, 12.5 ng/ml, 25 ng/ml, and 50 ng/ml), p-value calculated using one way ANOVA. Number of colonies are 1024, 1102 1134, and 1135 for the respective conditions. Results pooled from two separate experiments. Data represented as mean (s.d). G) Representative immunofluorescence images of SOX2, BRA, and CDX2 expression in geometrically confined 250µm diameter colonies differentiated in 6.25 ng/ml, 12.5 ng/ml, 25 ng/ml and 50 ng/ml of BMP4. Scale bars represent 200µm.

Taken together, these data are consistent with our hypothesis that the fate acquisition in the geometrically confined hPSC colonies, in response to BMP4, arises via a coordinated RD and PI model for cell fate patterning (**Fig 7**).

**Figure 7:**
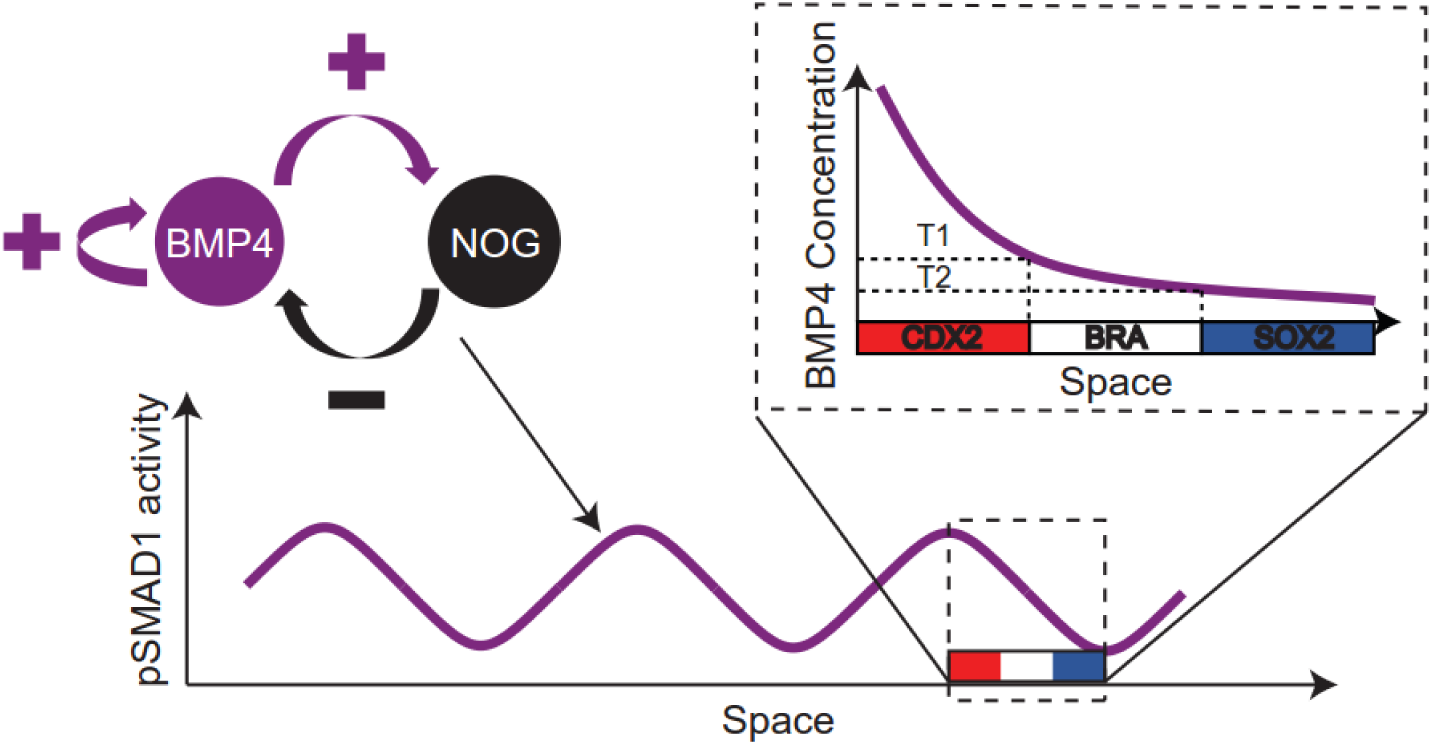
Mechanism of peri-gastrulation-like fate patterning in geometrically confined hPSC colonies. A BMP4-NOGGIN RD network induces a radial and periodic pSMAD1 activity gradient in geometrically confined hPSC colonies. The fate acquisition due to the signaling gradient proceeds following the classical PI fate patterning paradigm.

## Discussion

Of the various models of biological pattern formation that have been proposed, RD and PI have emerged as the dominantly accepted mechanisms to describe tissue organization during early development. However, due to apparent inconsistencies between the two models, RD and PI have historically been considered mutually exclusive. For instance, the RD hypothesis aimed to explain the mechanism of symmetry breaking during development^23,24^ whereas PI explicitly relied on the prior presence of polarities and asymmetries in the developing tissue^26,29^. Furthermore, the RD model implies a very close and bijective correspondence of the patterned fates with the morphogen distribution^23,24,34^ whereas PI requires a sufficiently graded distribution to pattern multiple fates, needing a stage of interpretation of the morphogen concertation^22,25,26,29^. Consequently, fate acquisition during development have predominantly been attributed to either RD^35–38^ or PI^39–44^ mechanisms. Although biological fate patterning occurring by both RD and PI working in concert has recently been hypothesized^22^, little evidence of this idea has been demonstrated. Our study provides evidence of RD mediated self-organization of a morphogen gradient (reported by pSMAD1 spatial patterning) which proceeds to pattern fates in a time- and dose-dependent manner consistent with the PI paradigm.

Etoc *et al*.^21^ proposed that BMP4 induced fate patterning in hPSC colonies is regulated by a negative feedback on BMP4 activity mediated by both a BMP-signaling induced NOGGIN expression, and reduced sensitivity to BMP ligands at the center of the colony. Although their model does predict the formation of a radial signaling gradient, it does not describe a classical RD system as it lacks a positive feedback of the activator^23,24,34^, and therefore, would never give rise to periodic RD-like patterns (**Sup Fig 14**). In contrast, we demonstrated that the presence of BMP4 in our cultures elicits both a positive feedback response as well as a negative feedback response mediated by the upregulation of NOGGIN, regardless of the basal medium used. BMP4 and NOGGIN form a stereotypical activator-inhibitor pair as per the RD paradigm, as BMP4 can induce short range activation of BMP signaling^32^, while NOGGIN is a highly-diffusible molecule that can result in long-range inhibition of BMP signaling^45,46^. Importantly, we explicitly demonstrate the induction of an RD-like response in larger colonies upon high dose induction of BMP4, both in the pSMAD1 response at 24 hours after induction and the fate acquisition 48 hours after induction.

The conventional understanding of the RD paradigm permits the short-range activation, and long-range inhibition is mediated by differential diffusivities of the activator and the inhibitor molecules – specifically, the diffusivity of the inhibitor is assumed to be greater than that of the activator^23,24,34^. This idea has even been specifically demonstrated to be true in the case of the NODAL, and LEFTY mediated formation of the Nodal signaling gradient during zebrafish embryogenesis^36^. Interestingly, our data show that NOGGIN (inhibitor) expression precedes that of BMP4 (activator) expression. A transcription factor network downstream of BMP signaling which results in a temporal lag between the expression of NOGGIN and BMP4 would enable greater long range accumulation of NOGGIN without requiring differential diffusivities of the molecules. Although the dynamics of the mechanism can still be captured very elegantly with the equations that were provided by Alan Turing^23^, our data hints at alternate mechanisms that can achieve RD patterns without requiring differential diffusivities of the activator and inhibitor molecules^47,48^.

We specifically tested and demonstrated that consistent with the PI paradigm, fate acquisition in the differentiating hPSC colonies in response to the pSMAD1 gradient arise both as a function of pSMAD1 concentration and duration of induction. Notably however, although the fate acquisition mediated by the pSMAD1 gradient follows the PI paradigm, the primitive-streak-like region is not specified in the absence of Nodal signaling. This underscores that coordination of multiple signaling pathways is required to pattern developing tissues. This is reminiscent of established models of PI mediated fate patterning like the gap genes in the *Drosophila* – which arise via a collaborative stimulation of multiple signals like Bicoid, Caudal, Hunchback etc^29^. Consequently, we propose that although pSMAD1 activity patterns fates via the PI paradigm, it does so in conjunction with other transcription factors like SMAD2, the effector of Nodal signaling.

Morphogen gradients formed in developing embryos are thought to scale with size. For instance, the scaling of the dorsal-ventral axis in the developing *Xenopus* is robust to dramatic manipulations like resection of the ventral half – in which case, the embryo is able to develop into a smaller but proportionally patterned larva^49^. This characteristic had been attributed to the function of the Spemann’s organizer^46,49^. This underscores the fact that the robustness of the induction and stabilization of the morphogen gradients that pattern a developing embryo is the result of a concerted effort by multiple tissues. Our data show that the fate organization in the differentiating micro-patterned hPSC colonies of varying sizes is not robust to a specific BMP4 concentration. Notably however, modulation of the BMP4 dose in the induction media can recapitulate the appropriate fate patterning. This highlights that robustness to changes in the size of the developing epiblast is attainable, and emphasizes the importance of regulation from other tissues like the vertebrate organizer (the primitive node) in conferring robustness to a developing human embryo. The mechanism that regulates the pattern formation in the micro-patterned hPSC colonies undergoing peri-gastrulation-like events is one of the multiple layers of complexity that would be present in a developing human embryo.

In conclusion, we report a defined, *in vitro* model of human peri-gastrulation-like biological fate patterning, and demonstrate that the mechanistic underpinning of this observation is a step-wise model of both RD and PI. In doing so we provide evidence that these two models can work in a coordinated manner. Our data indicates possible alternate mechanisms of enabling short range activation and long range inhibition, characteristic of the RD model, independent of requiring the inhibitor diffusivity to be higher than that of the activator. We further report that fate acquisition specific to human peri-gastrulation-like events occur in a manner consistent with the PI paradigm and require multiple signaling pathways working in concert. Finally, our data implicates the need of multiple tissues cooperating to induce morphogen gradients that can scale and maintain robustness to perturbations that may occur in the developing embryo. Consequently, our work not only provides deep insight into one of the earliest stages of human embryonic development, but also into the general mechanisms employed in patterning of biological form.

## Materials and Methods

### Human Pluripotent Stem Cell Culture

CA1 hPSC line (generously provided by Andras Nagy, Samuel Lunenfeld Research Institute) were cultured on Geltrex (diluted 1:50) coated 6-well tissue culture plates using mTeSR1^TM^ medium (StemCell Technologies) as per manufacturer’s instructions. The cells were passaged at a ratio of 1:12 using ReleSR^TM^ (StemCell Technologies) per manufacturer’s instructions. For the first 24 hours after passage, the cells were cultured in mTeSR1 supplemented with ROCK inhibitor Y-27632 to increase cell viability. The medium was changed every day and passaged every 4-5 days or when the cells were 75-80% confluent. The cell line was tested for contamination.

### Preparation of PEG plates to micro-pattern hPSC colonies

Custom sized (110mmx74mm) Nexterion-D Borosilicate thin glass coverslips (SCHOTT) were activated in a plasma cleaner (Herrick Plasma) for 3 minutes at 700 mTorr, and incubated with 1 ml of Poly-L-Lysine grafted Polyethylene Glycol (PLL-g-PEG, SUSOS, cat # PLL(20)-g(3.5)-PEG(5)) at a concentration of 1 mg/ml at 37°C overnight. The glass slides were then extensively rinsed with ddH_2_O and dried. The desired patterns were transferred to the surface of the PEG-coated side of the coverslip by photo-oxidizing select regions of the substrate using Deep UV exposure for 10 minutes through a Quartz photomask in a UV-Ozone cleaner (Jelight). Bottomless 96-well plates were plasma treated for 3 minutes at 700 mTorr and the patterned slides were glued to the bottomless plates to produce micro-titer plates with patterned cell culture surfaces. Prior to seeding cells onto the plates, the wells were activated with N-(3-Dimethylaminopropyl)-N′-ethylcarbodiimide hydrochloride (Sigma cat# 03450), and N-Hydroxysuccinimide (Sigma, cat# 130672) for 20 minutes. The plate was thoroughly washed three times with ddH_2_O, and incubated with Geltrex (diluted 1:150) for four hours at room temperature on an orbital shaker. After incubation, the plate was washed with Phosphate Buffered Saline (PBS) at least three times to get rid of any passively adsorbed extracellular matrix (ECM) and seeded with cells to develop micro-patterned hPSC colonies.

### Cell seeding and induction of peri-gastrulation-like events

To seed cells onto ECM-immobilized PEG-UV 96-well plates, a single cells suspension of the CA1 line was generated by incubation in 1ml of TryplE (Invitrogen) per well for 3 minutes at 37°C. The TryplE was blocked using in equal volume DMEM + 20% KnockOut Serum Replacement (SR) (Invitrogen) and the cells were dissociated by pipetting to generate a single cell suspension. The cells were centrifuged and re-suspended at a concentration of 1 × 10^6^ cells/ml in SR medium supplemented with 20ng/ml bFGF (R&D) and 10µM ROCK inhibitor Y-27632. SR medium consists of 74% DMEM, 1% Penicillin/Streptomycin, 1% non-essential amino acids, 0.1mM β-mercaptoethanol, 1% Glutamax, 2% B27 minus retinoic acid, and 20% SR (all Invitrogen). Wells were seeded in the PEG-patterned 96 well plates at a density of 80,000 cells/well and incubated for 2 hours at 37°C. After 2 hours, the medium was changed to SR without ROCKi. When confluent colonies were observed (12-16 hours after seeding), the peri-gastrulation-like induction was initiated in N2B27 medium supplemented with BMP4 (R&D) (BMP4 dose depended on experimental design). The peri-gastrulation-like pattern formation assay was typically performed with 1000 µm diameter colonies 48 hours following induction, although additional time points and colonies sizes were tested as described in the Results section for specific experiments. N2B27 medium consists of 93% DMEM, 1% Penicillin/Streptomycin, 1% non-essential amino acids, 0.1mM β-mercaptoethanol, 1% Glutamax, 1% N2 Supplement, 2% B27-retinoic acid supplement (all Invitrogen) supplemented with 100 ng/ml Nodal (R&D), and 10 ng/ml bFGF (R&D).

### Single cell data acquisition and analysis of immunofluorescence imaging

The patterned plates were fixed with 3.7% paraformaldehyde for 20 minutes, rinsed three times with PBS and then permeabilized with 100% methanol for 3 minutes. After permeabilization, the patterned colonies were blocked using 10% Fetal Bovine Serum (Invitrogen) in PBS overnight at 4ºC. Primary antibodies were incubated at 4ºC overnight. The following day the primary antibodies were removed, and the plates were washed three times with PBS followed by incubation with the secondary antibodies and DAPI nuclear antibody at room temperature for 1 hour. Single cell data was acquired by scanning the plates using the Cellomics Arrayscan VTI platform using the TargetActivation. V4 bioassay algorithm. This algorithm utilizes the expression intensity in the DAPI channel to identify individual nuclei in all fields imaged and acquires the associated intensity of proteins of interest localized within the identified region. The single cell data was exported into Context Explorer (CE), a custom software developed in-house for image analysis (Östblom, *et al*., unpublished). In CE, cell colonies are identified through the DBSCAN algorithm as implemented in Python’s Scikit-learn package. Within a colony, each cell is assigned x- and y-coordinates relative to the colony centroid. To create the colony overlay plots, cells from multiple colonies are grouped in hexagonal bins per their relative x- and y-coordinates. These positional bins are color-coded to represent the average protein expression level of all cells within a bin. The color map range is normalized to the lowest and highest expressing hexagonal bins. Spatial expression trends within colonies are also visualized as line plots, where cells are grouped by the Euclidean distance between a cell and the centroid of the colony. For each colony, the average expression value of all cells within a distance bin is computed. The line plots describe the mean expression value, standard deviation and 95% confidence interval (CI) between colonies as indicated in figure legends. The line plots depicting radial trends of proteins of interest in individual colonies were acquired through the sections of the colonies depicted (shown as white rectangles in the figures) in Fiji (ImageJ). The plot profile extracted was then run through a Savitzky-Golay smoothing filter in Matlab and represented as a function of radial distance.

**Table 1:** Antibodies used for immunofluorescence experiments

|  |  |
| --- | --- |
| CDX2 | Abcam (ab15258, ab76541) |
| BRA | R&D (AF2085) |
| SOX2 | Cell Signaling (3579S), R&D (MAB2018) |
| SOX17 | R&D (AF1924) |
| EOMES | Abcam (ab23345) |
| EPCAM | R&D (SC026) Kit |
| SNAIL | R&D (SC026) Kit |
| pSMAD1 | Cell Signaling (9516S) |

### Quantitative PCR analysis

RNA extraction was performed using Qiagen RNAeasy miniprep columns according to the manufacturer’s protocol, and the cDNA was generated using Superscript III reverse transcriptase (Invitrogen) according to the manufacturer’s instructions. The generated cDNA was mixed with primers for the genes of interest and SYBR green mix (Roche, Sigma) and the samples were run on an Applied Biosystems QuantStudio 6 flex real-time PCR machine. Relative expression of described genes was determined by the delta–delta cycle threshold (ΔΔCt) method with the expression of GAPDH as an internal reference. Primer sequences used are provided in Supplementary Information.

**Table 2:**
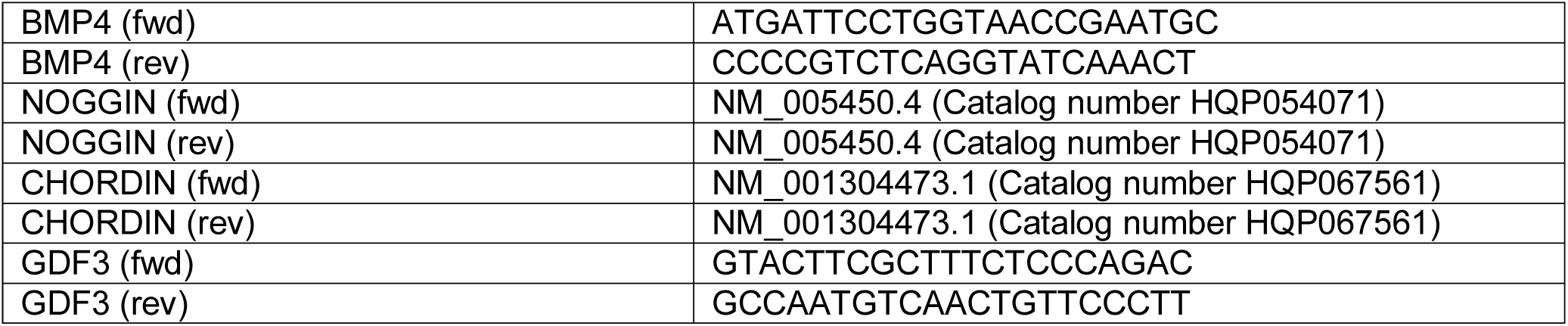
Primers used in qPCR experiments

### Statistics and data analysis

All gene expression results were expressed as mean (+S.D.). Statistical tests for gene expression results were performed using two-tailed Student’s t-test assuming unequal variance between datasets. The statistical significance for fate acquisition results were calculated using one-way ANOVA on ranks (Kruskal-Wallis test). The calculations for Student’s t-tests were performed in Excel, and the one-way ANOVA calculations were performed in Prism.

## Acknowledgements

We thank Dr. Andras Nagy for kindly donating the CA1 human embryonic stem cell lines. We thank Dr. Janet Rossant, Dr. Celine Bauwens, and Dr. Geoff Clarke for providing insightful feedback on our manuscript. We thank Dr. Aryeh Warmflash for helpful correspondence. We would like to acknowledge CMC microsystems for the provision of products and services including Comsol. This work was funded by the the Canadian Institutes for Health Research (CIHR) and Medicine by Design, a Canada First Research Excellence Program at the University of Toronto. M. T. received funding from the CREATE M3 from the Natural Sciences and Engineering Research Council of Canada. PWZ is the Canada Research Chair in Stem Cell Engineering.

## Supplemental figure legends

**Sup Figure S1:**
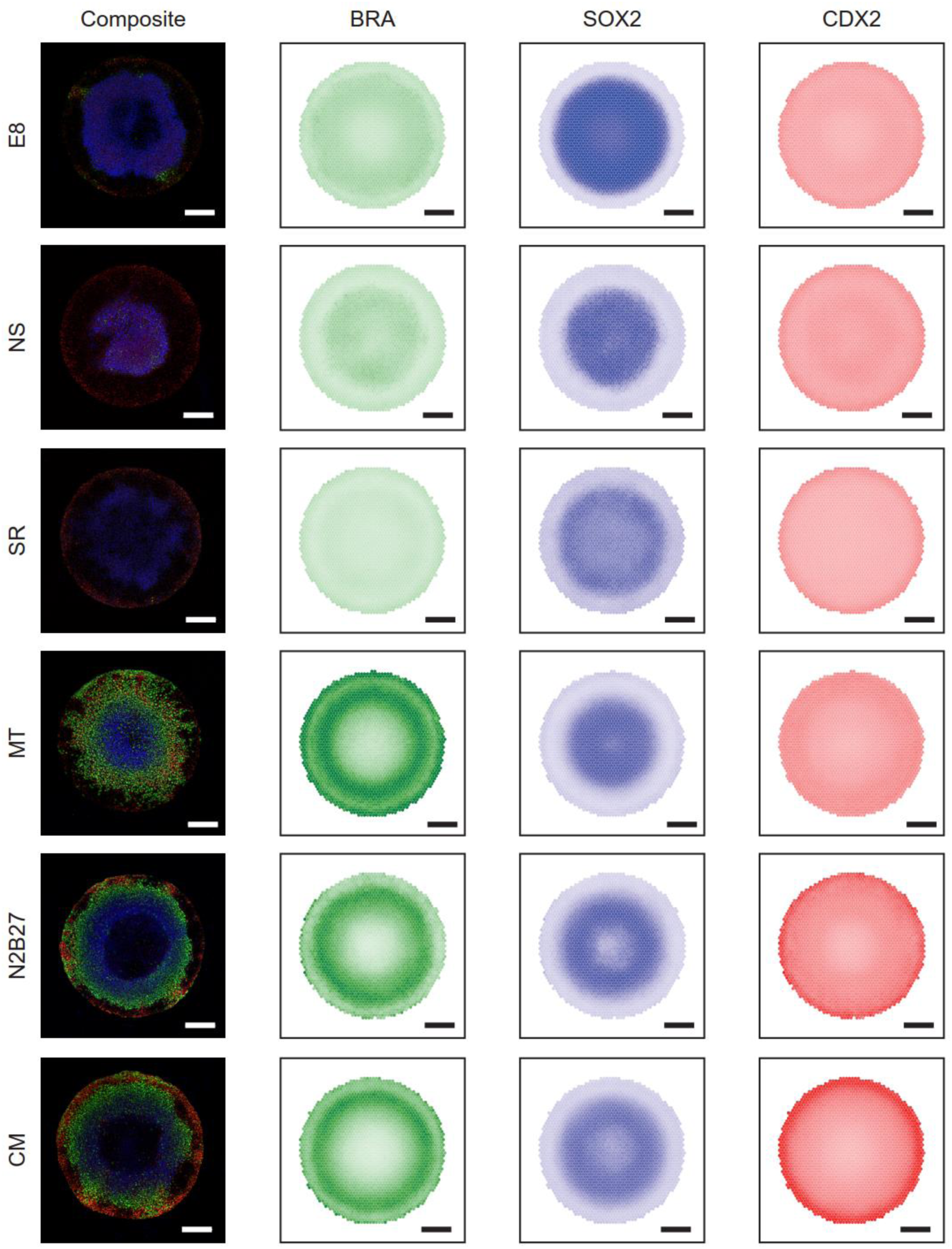
Peri-gastrulation-like fate patterning in multiple basal medium conditions. A) Representative composite images of SOX2, BRA and CDX2 staining in micro-patterned hPSC colonies differentiated in BMP4 supplemented E8, Nutristem (NS), SR, mTeSR, and N2B27 (left column), and spatial expression overlays for each specific fate marker performed in CE (3 right panels). Scale bars represent 200µm.

**Sup Figure S2:**
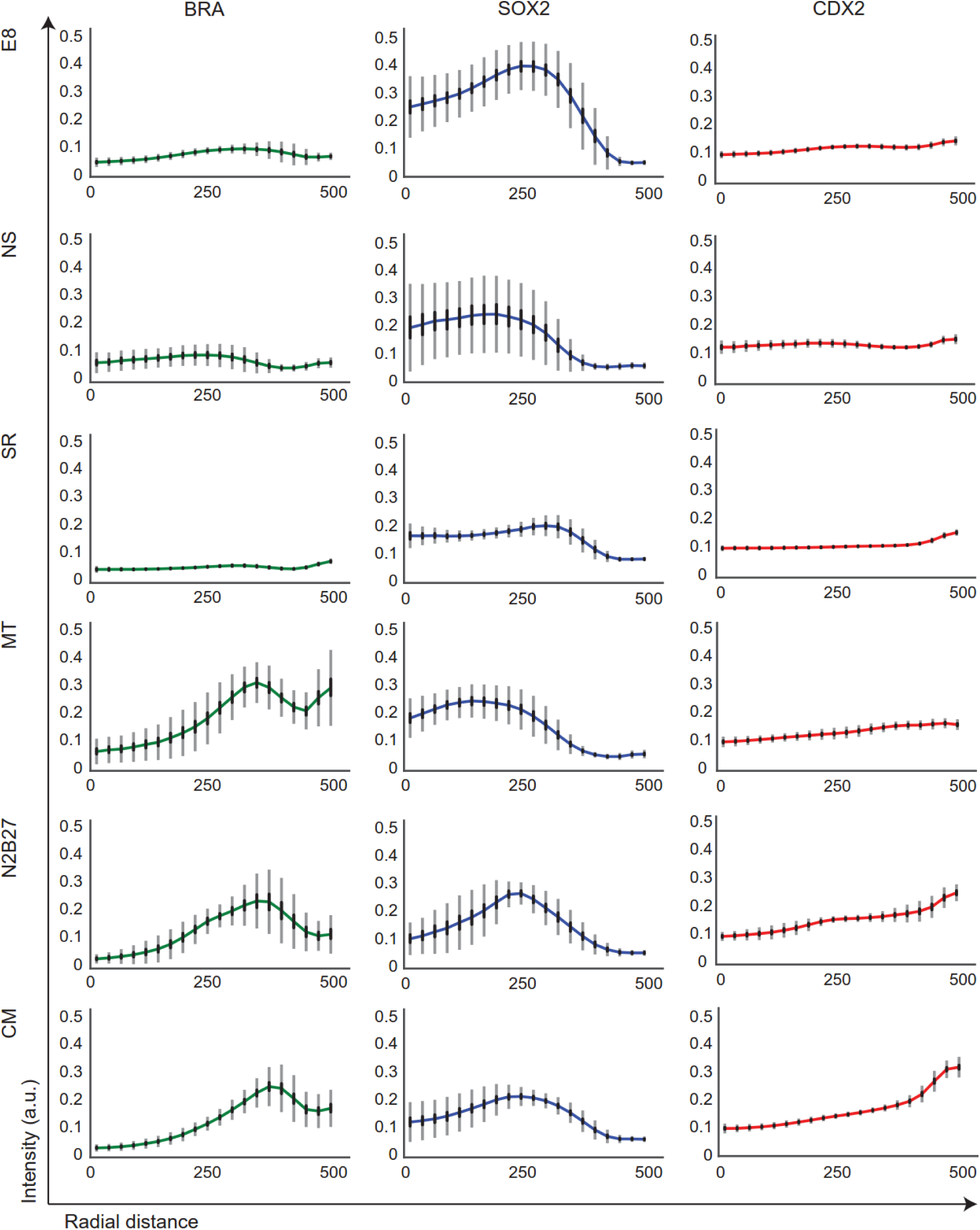
Spatial trends of CDX2, BRA, and SOX2 observed in different basal media. Radial trends of A) CDX2, B) BRA, and C) SOX2 in BMP4 supplemented E8, NS, SR, MT, N2B27, and CM. Standard deviations shown in grey, and 95% confidence intervals shown in black.

**Sup Figure S3:**
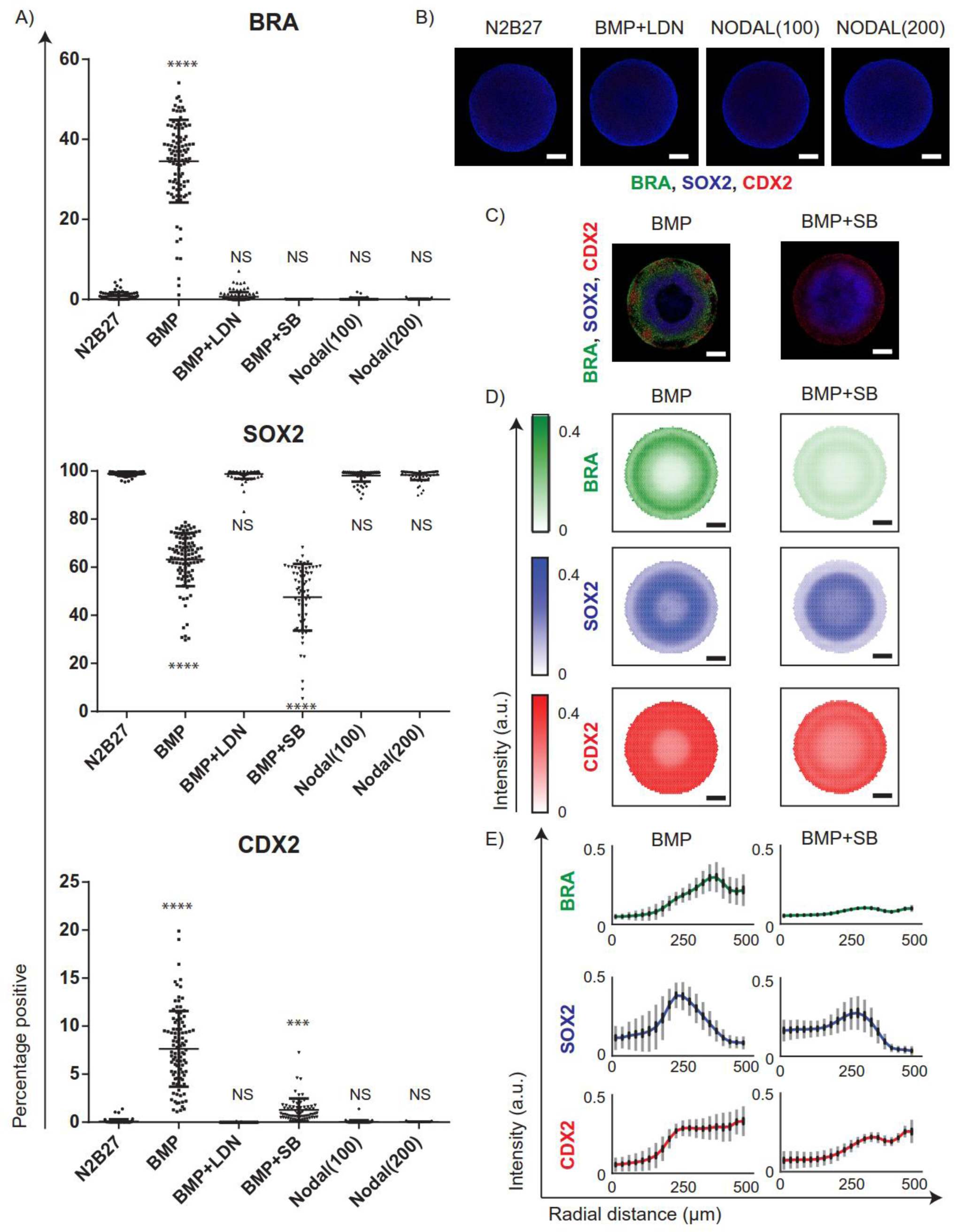
Nodal signaling is required for primitive streak specification, but does not induce differentiation and fate patterning in micro-patterned hPSC colonies. A) Percentage of cells expressing BRA, SOX2, and CDX2 in N2B27 (n=73), N2B27+BMP (n=100), N2B27+BMP+LDN (n=101), N2B27+BMP+SB (n=67), N2B27+100ng/ml Nodal (n=68), and N2B27+ 200ng/ml Nodal (n=65). The experiment was performed twice. **** (p<0.0001), NS (p>0.05). The p values were calculated using one-way ANOVA. B) Representative images of colonies cultured in N2B27, BMP+LDN, Nodal (100), and Nodal (200) stained for BRA, SOX2, and CDX2. C) Representative immunofluorescent images of colonies cultured in BMP4, and BMP4+SB stained for BRA, SOX2, and CDX2. D) Overlaid SOX2, BRA and CDX2 expression averages for 100 colonies cultured in BMP4, and 67 colonies cultured in BMP4+SB. E) Average radial trends for BMP4 and BMP4+SB conditions. Standard deviations shown in grey, and 95% confidence intervals shown in black. Scale bars represent 200µm.

**Sup Figure S4:**
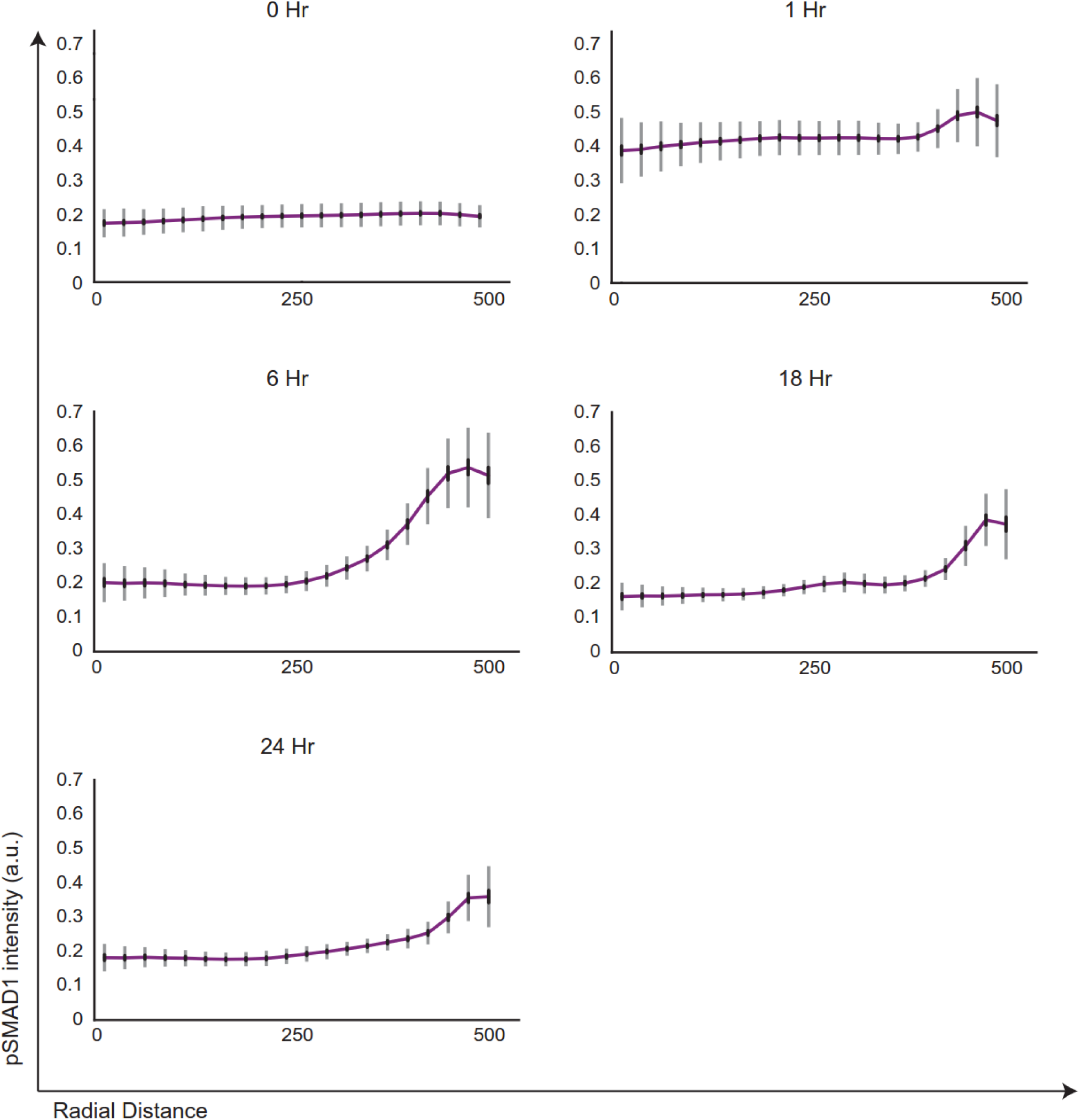
Quantified radial trends of pSMAD1 activity over the first 24 hours after BMP4 induction. Radial trends of pSMAD1 expression observed in BMPi = 50ng/ml 0, 1, 6, 18, and 24 hours following induction. Standard deviations shown in grey, and 95% confidence intervals shown in black. The number of colonies per condition are 202, 193, 105, 100, and 105 respectively pooled from two experiments.

**Sup Figure S5:**
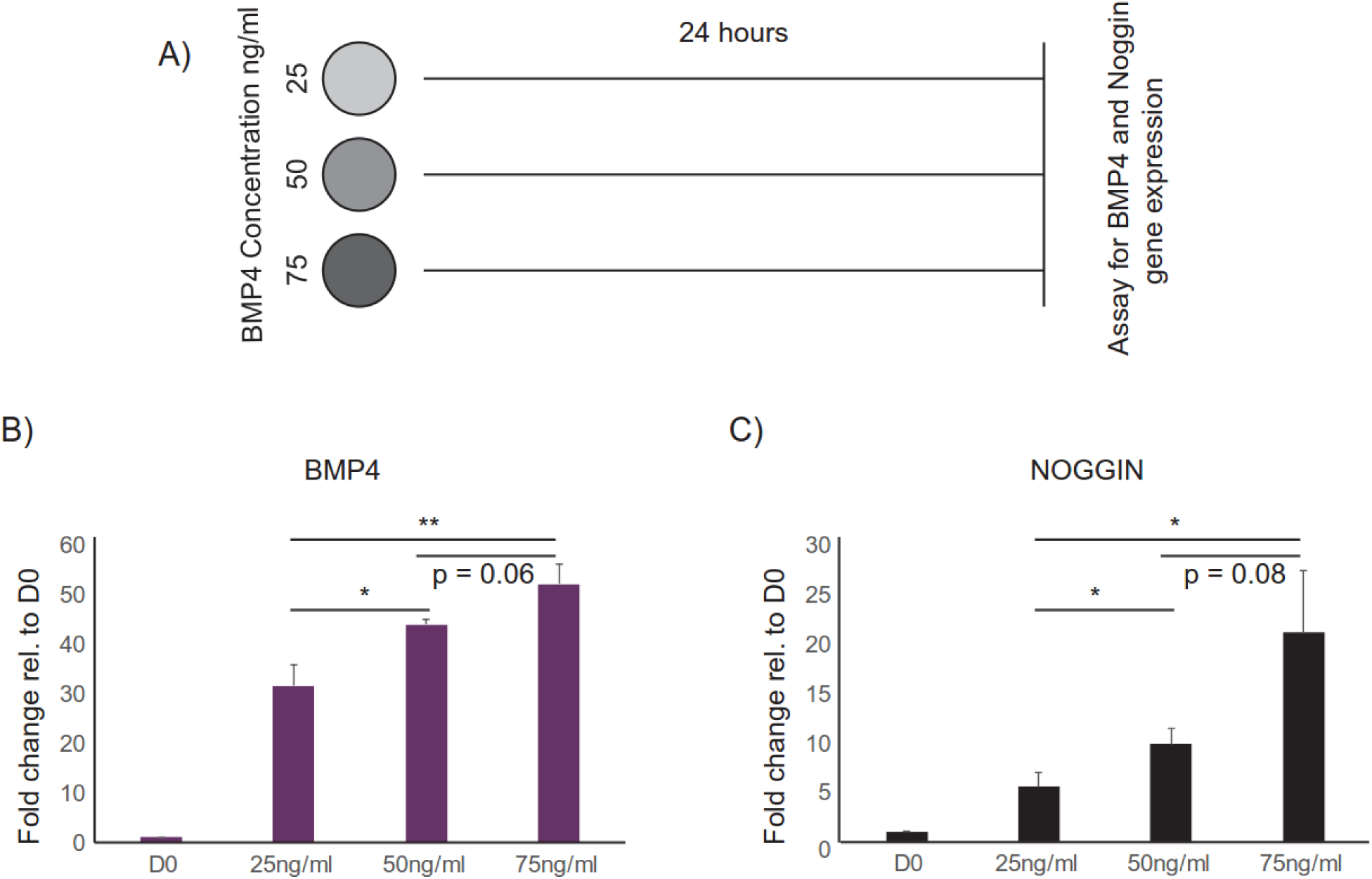
BMP4 and NOGGIN upregulation occur in a BMP4 induction dose-dependent manner. A) Experimental overview: Gene expression data gathered at 24 hours following induction at varying concentrations of BMP4. B) BMP4-induced expression of BMP4. C) BMP4 induced expression of NOGGIN. Data represented as mean and S.D. of three biological replicates. * p<0.05, ** p<0.01.

**Sup Figure S6:**
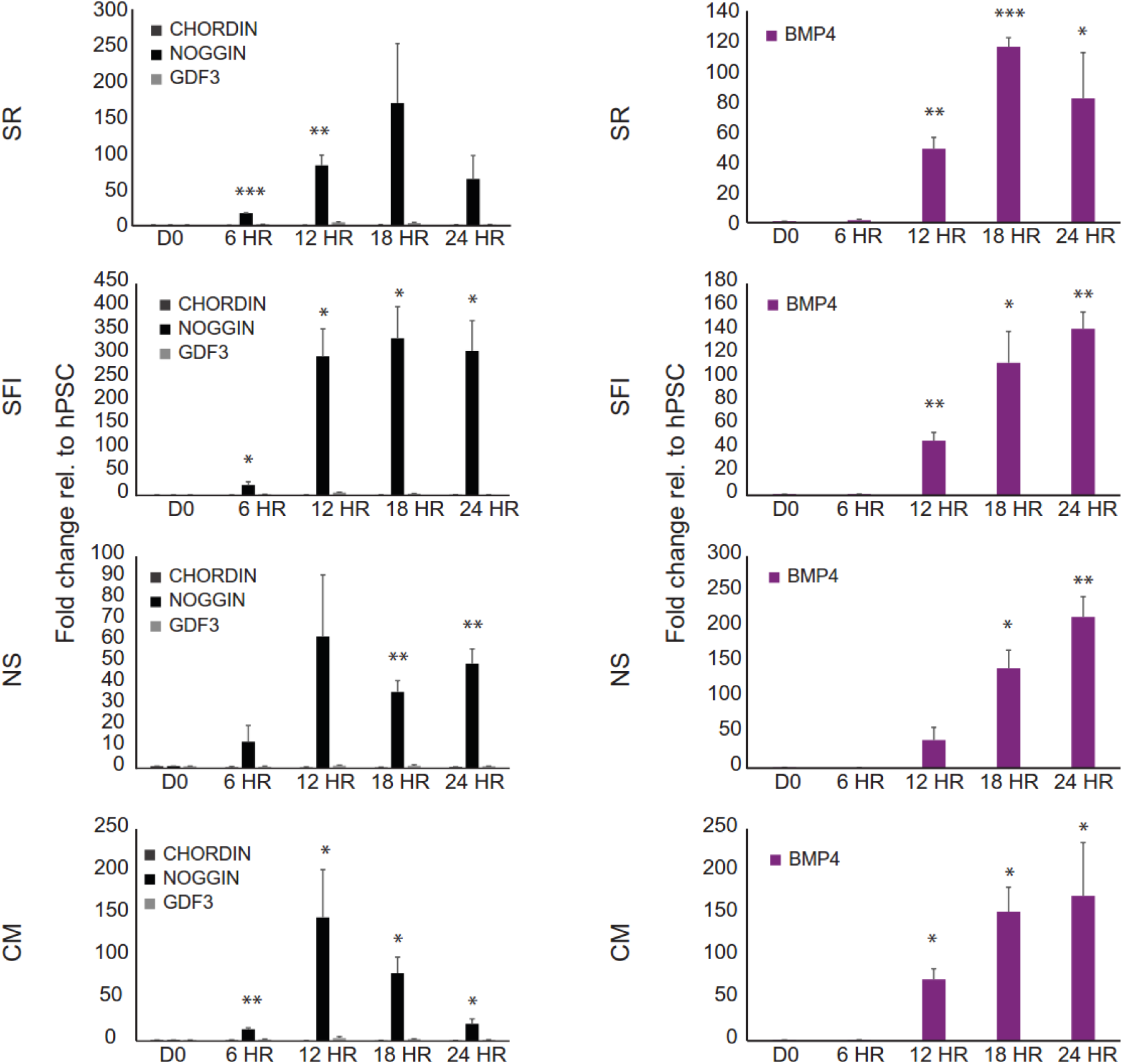
BMP4 induced upregulation of BMP4 and NOGGIN in tested medium conditions. A) Kinetic gene expression profiles for BMP4 and its cardinal inhibitors in response to BMP4 induced differentiation. Medium conditions tested include a Knockout serum – based medium (SR), a serum-free medium (SFI – for composition, please see Nazareth *et al*., Nature Methods 2013), Nutristem (NS), and Mouse Embryonic Fibroblast conditioned medium (CM). Data represented as mean and S.D of three biological replicates. * p<0.05, ** p<0.01, *** p<0.001.

**Sup Figure S7:**
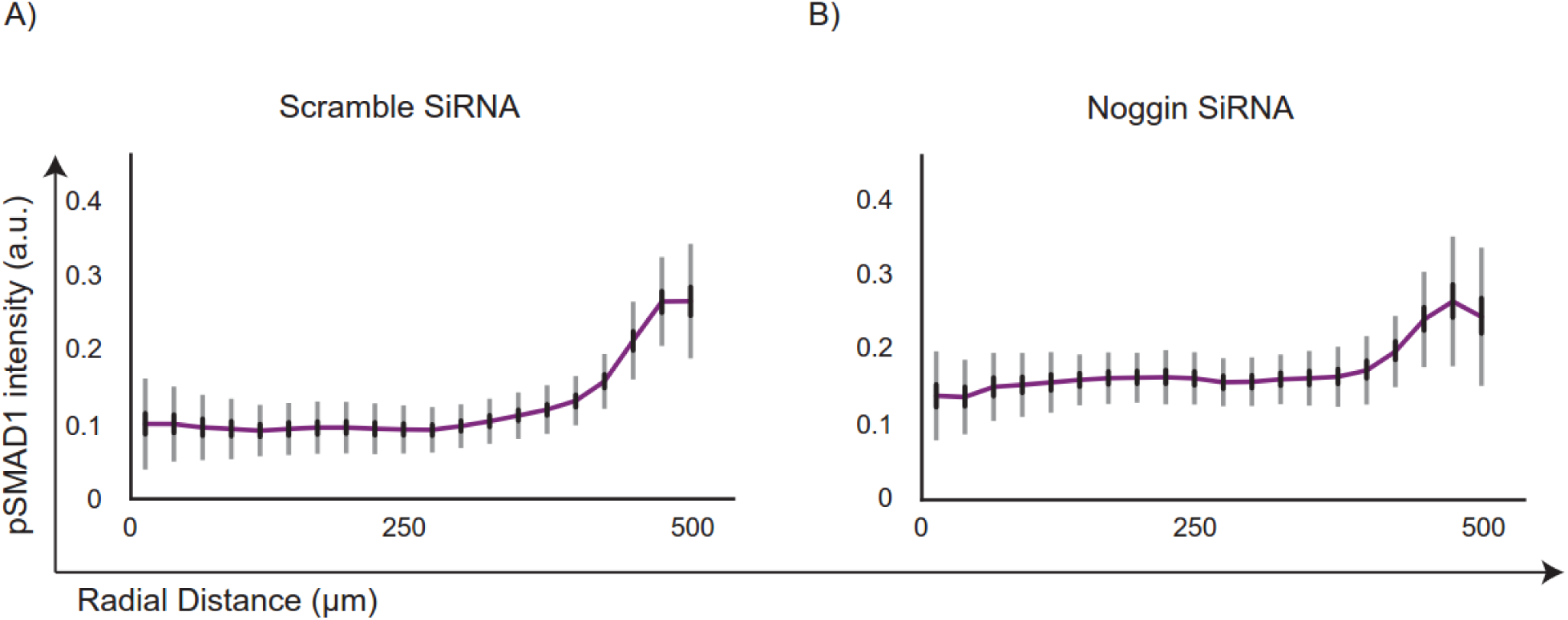
Quantified radial trends of pSMAD1 activity at 24 hours after NOGGIN inhibition using SiRNA. Radial trends of pSMAD1 observed in BMPi = 50ng/ml shown A) Scramble SiRNA (64 colonies), B) NOGGIN SiRNA (62 colonies). Standard deviations shown in grey, and 95% confidence intervals shown in black.

**Sup Figure S8:**
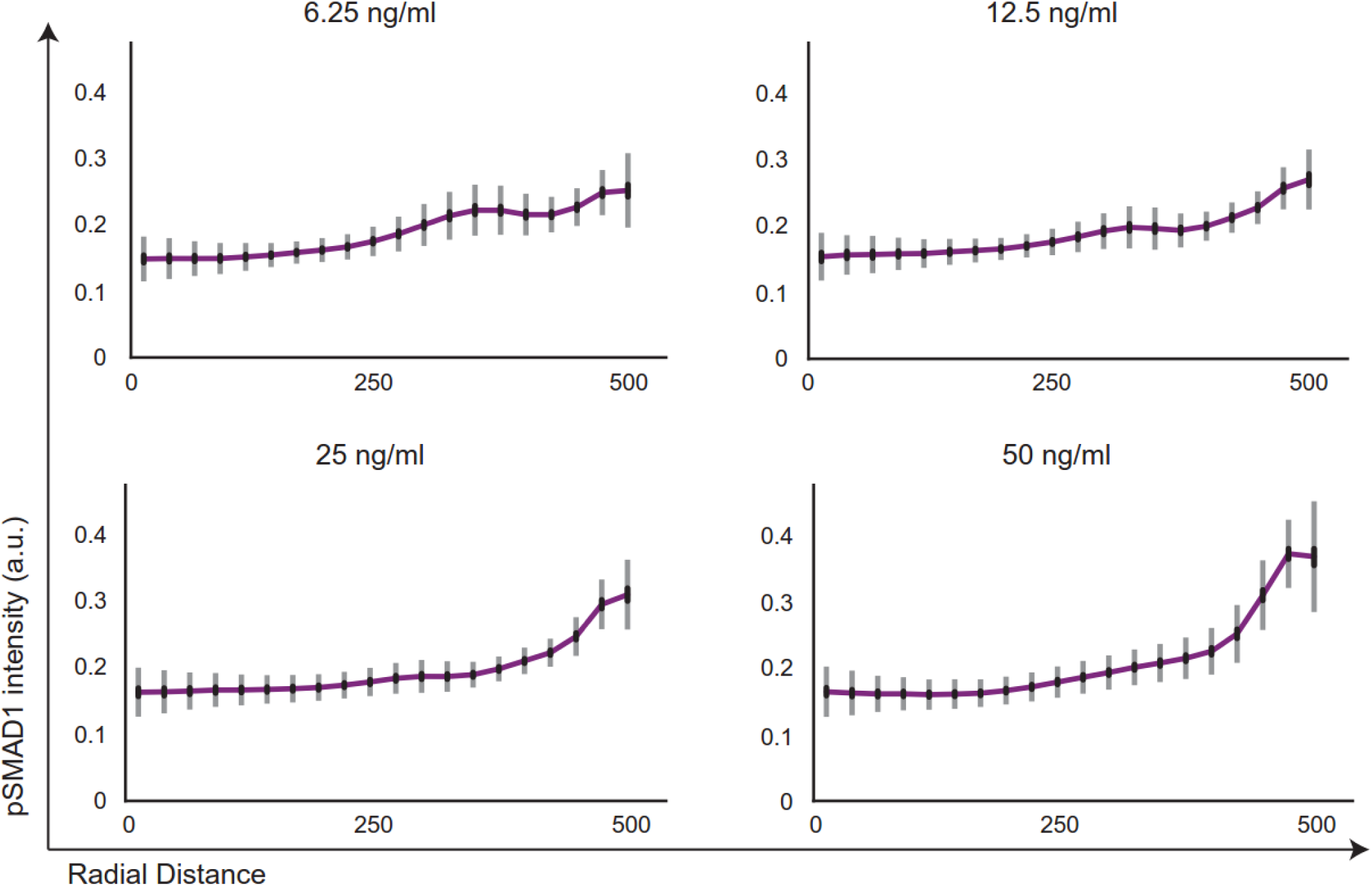
Quantified radial trends of pSMAD1 activity at 24 hours after induction with varying concentrations of BMP4. Radial trends of pSMAD1 activity were observed in varying BMPi concentrations (6.25 ng/ml, 12.5 ng/ml, 25 ng/ml, and 50ng/ml). Standard deviations shown in grey and 95% confidence intervals shown in black.

**Sup Figure S9:**
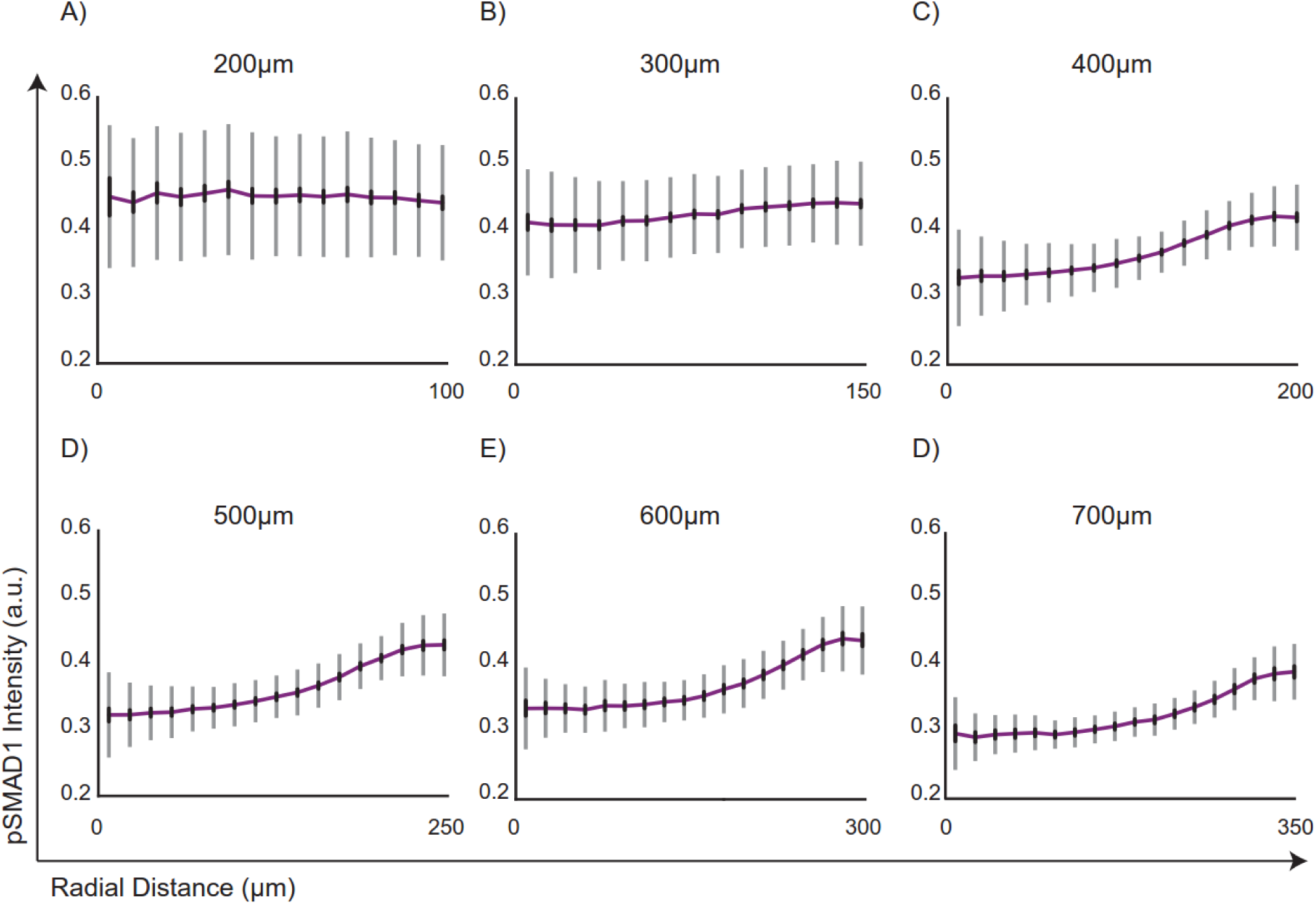
Quantified radial trends of pSMAD1 activity at 24 hours after BMP4 in colonies of varying sizes. Radial trends of pSMAD1 activity observed in BMPi = 50 ng/ml for varying colony sizes. The colony diameters range from A) 200 µm (373 colonies), B) 300 µm (437 colonies), C) 400 µm (261 colonies), D) 500 µm (178 colonies), E) 600 µm (118 colonies), and F) 700 µm (87 colonies). Standard deviations shown in grey, and 95% confidence intervals shown in black.

**Sup Figure S10:**
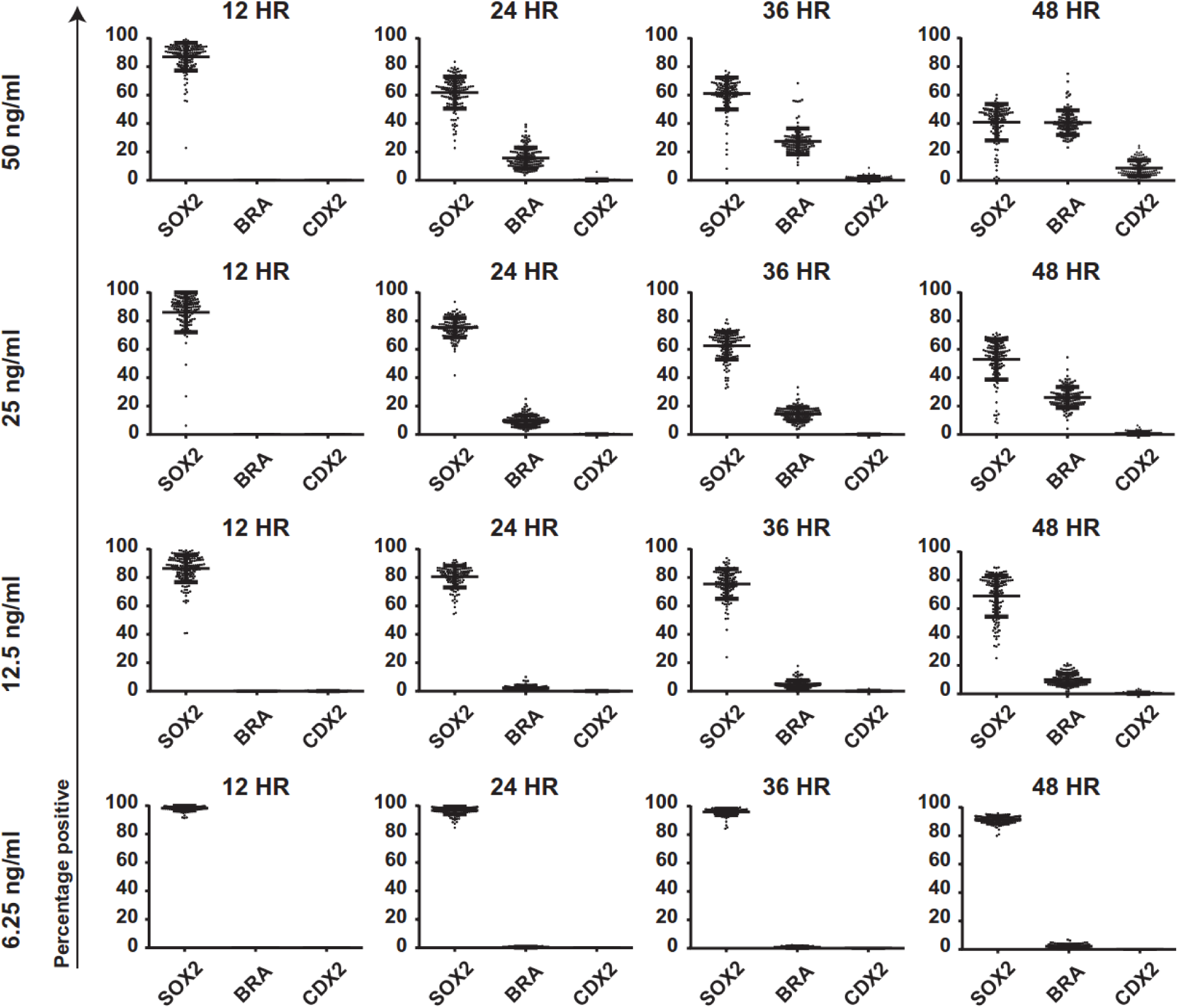
CDX2 and BRA expression in colonies arise as a function of BMP4 dose, and induction time. Percentage of cells expressing SOX2, BRA, and CDX2 in colonies induced to differentiate at varying concentrations of BMP4 (6.25 ng/ml, 12.5 ng/ml, 25 ng/ml, and 50 ng/ml) and induction times (12 hours, 24 hours, 36 hours, and 48 hours). Each condition had over 140 colonies. Data pooled from two experiments.

**Sup Figure S11:**
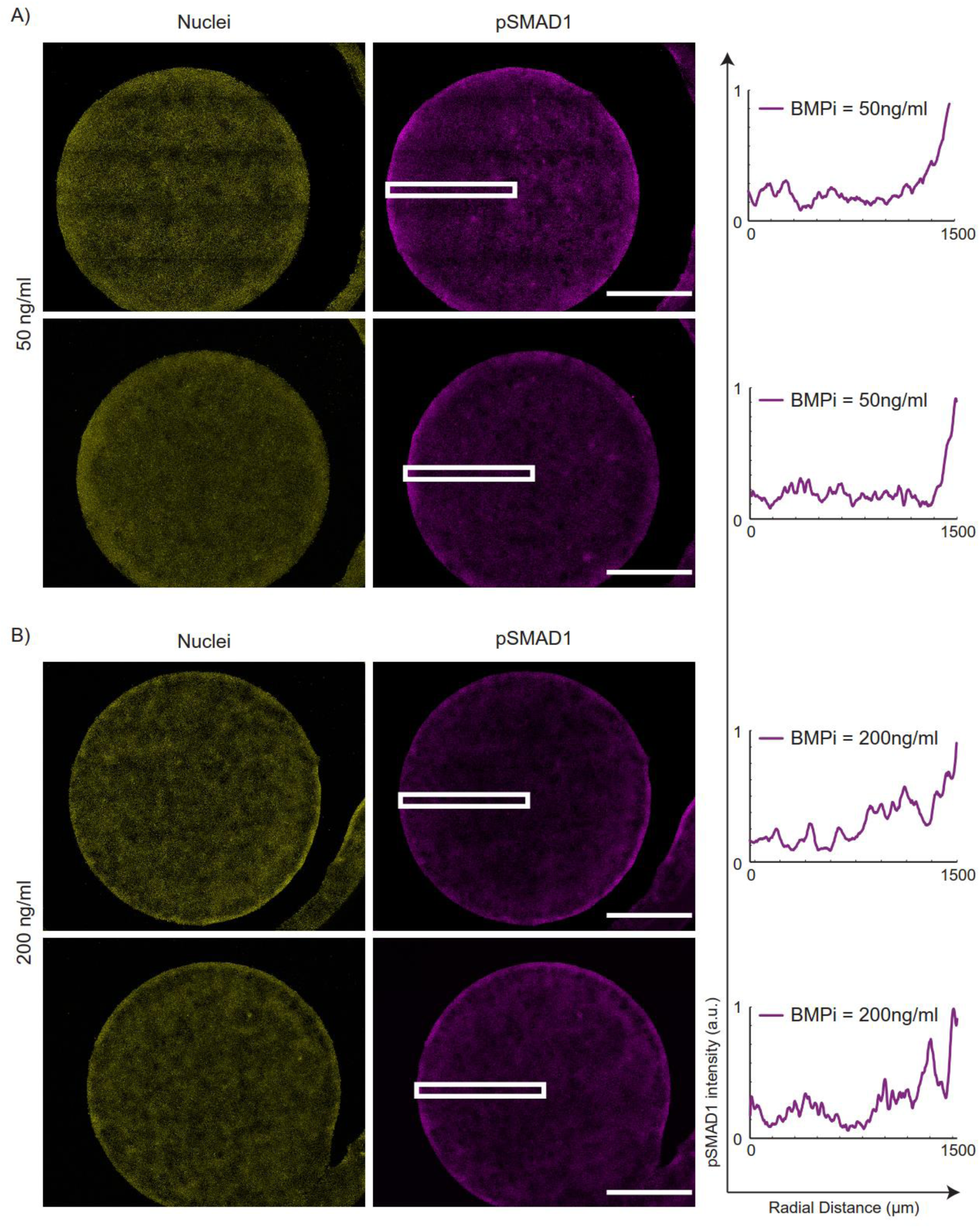
RD-like patterns noted in pSMAD1 activity in varying doses of BMP4. Representative immunofluorescent images of 3mm diameter colonies stained for DAPI, and pSMAD1 for a BMP4 dose of A) 50ng/ml, and B) 200ng/ml. Scale bars represent 1mm.

**Sup Figure S12:**
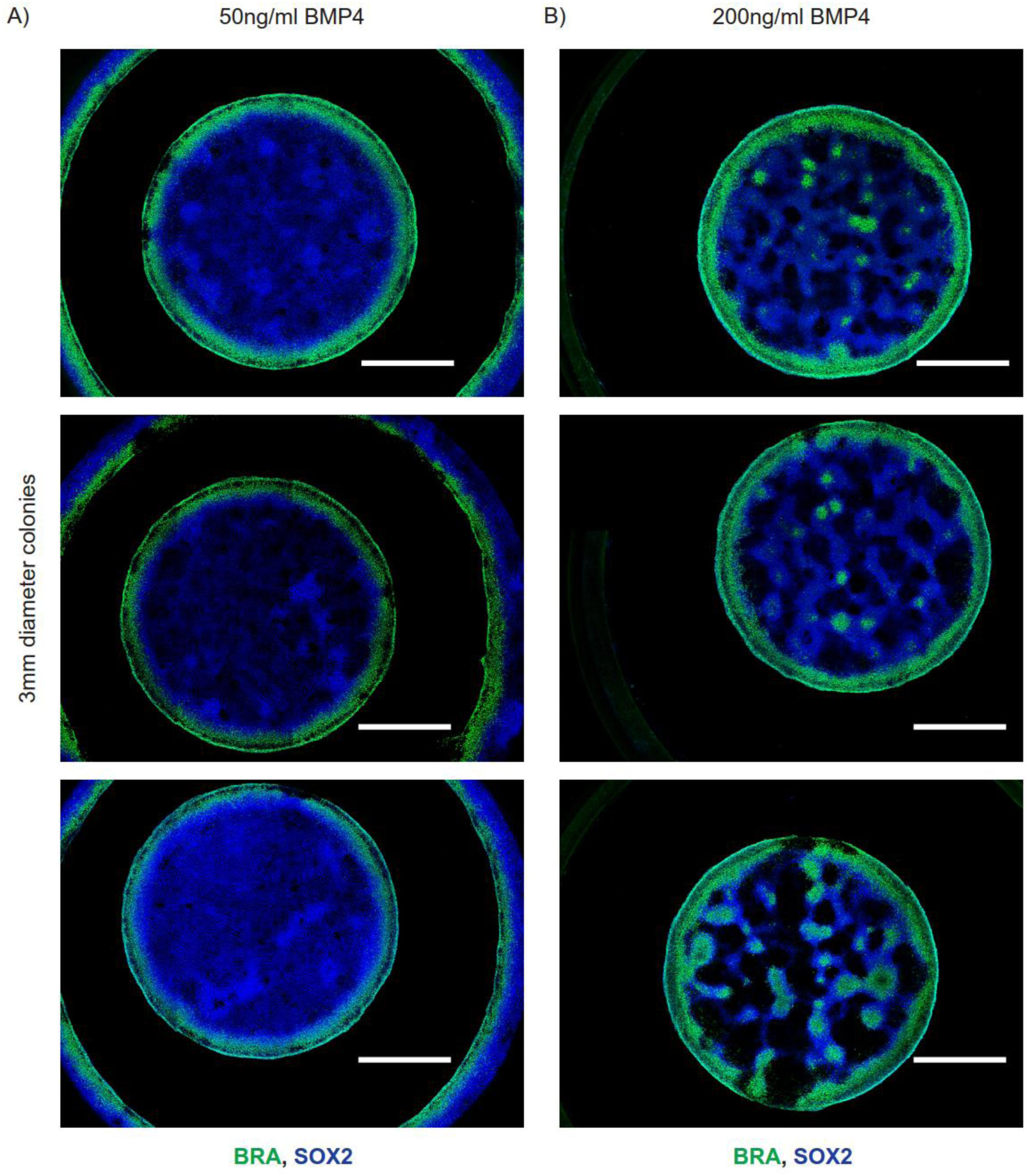
RD-like patterns noted in fate acquisition in varying doses of BMP4. Representative immunofluorescent images of 3mm diameter colonies stained for SOX2, and BRA for a BMP4 dose of A) 50ng/ml, and B) 200ng/ml. Scale bars represent 1mm.

**Sup Figure S13:**
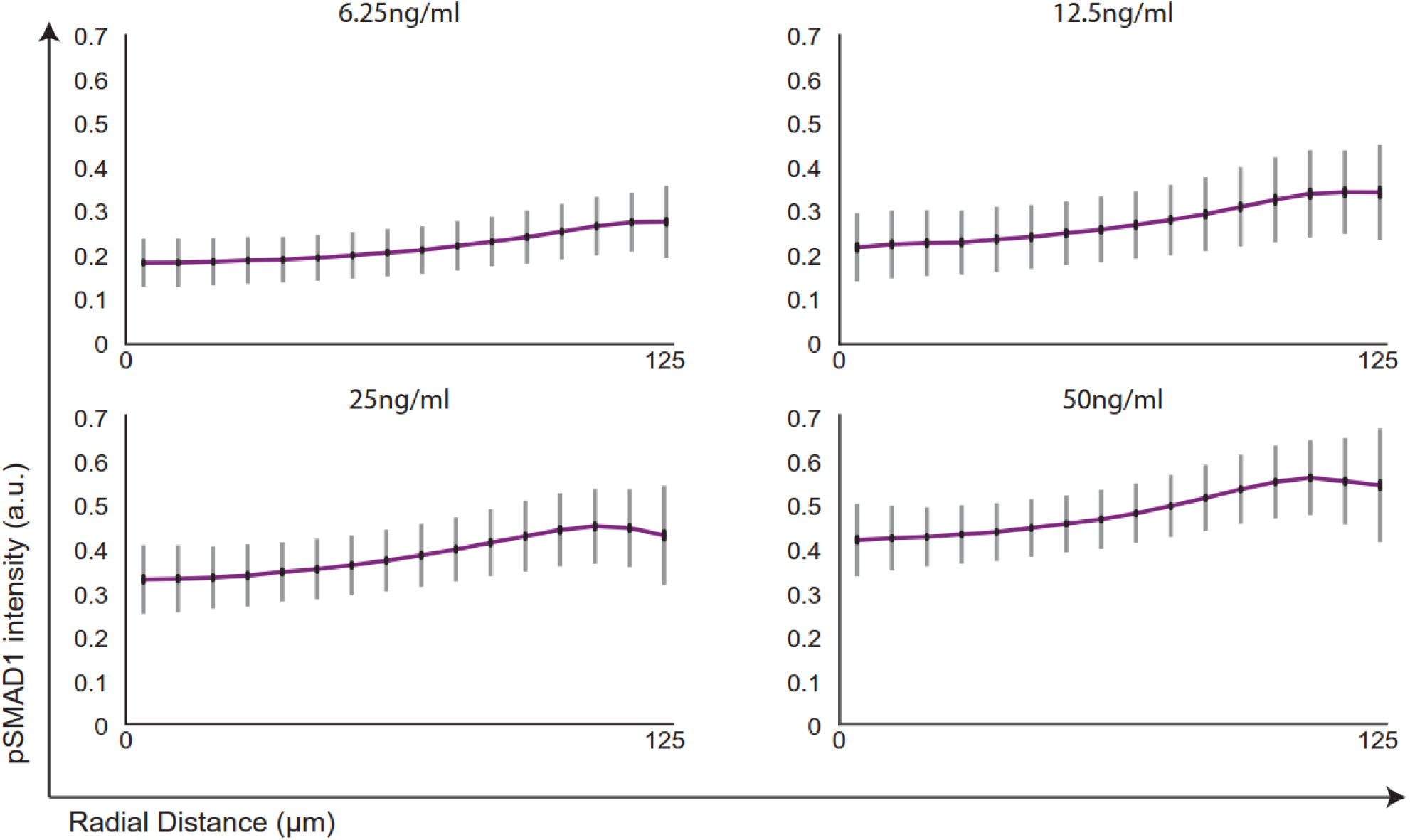
Quantified radial trends of pSMAD1 activity in colonies of 250µm diameter after 24 hours of BMP4 induction. Radial trends of pSMAD1 activity were observed in colonies of 250µm diameter 24 hours after induction with varying BMP4 concentrations – 6.25 ng/ml (976 colonies), 12.5ng/ml (638 colonies), 25ng/ml (475 colonies), and 50ng/ml (689 colonies). Standard deviations shown in grey and 95% CI shown in black.

**Sup Figure S14:**
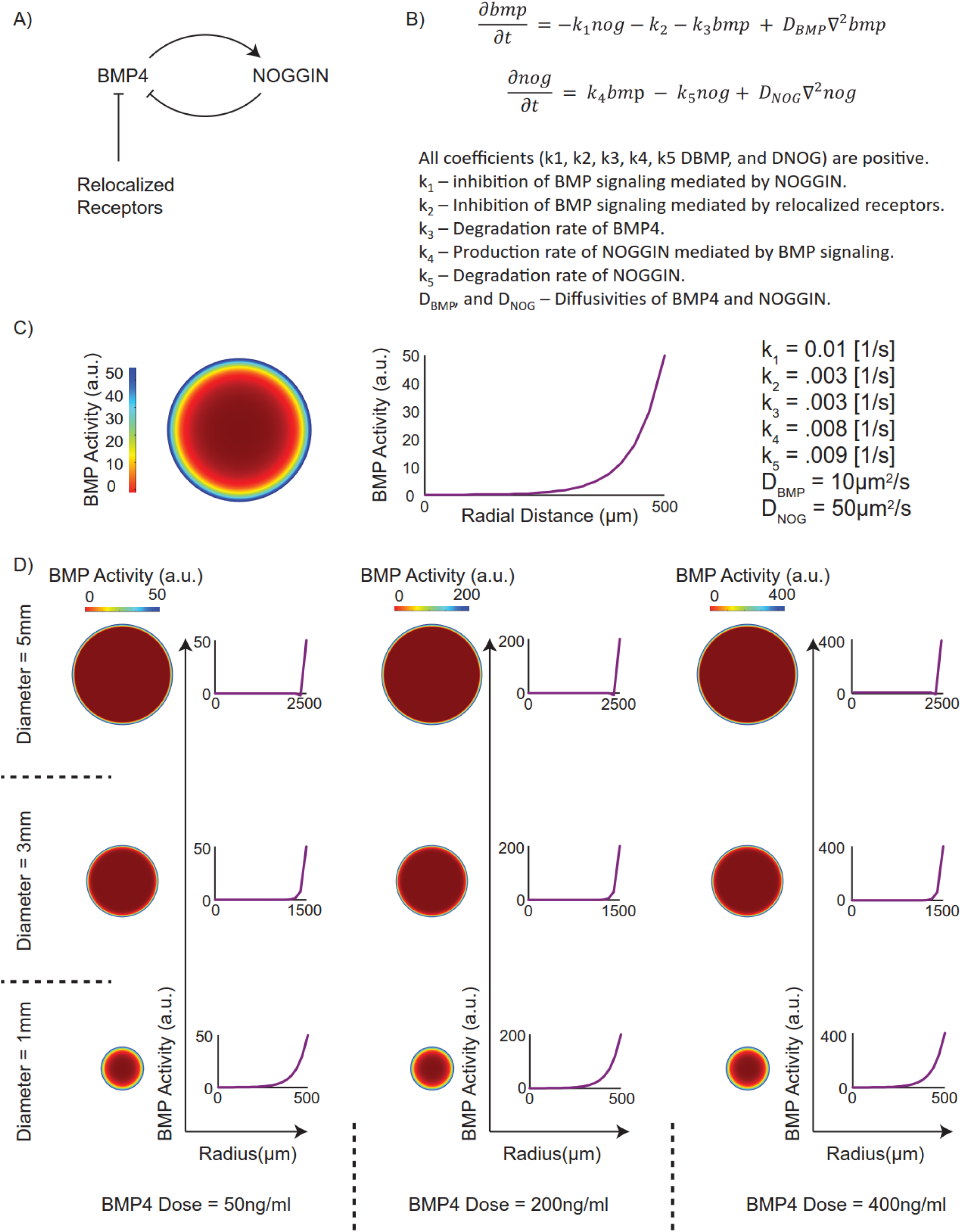
Dual inhibition model does not give rise to repetitive RD-like free BMP4 distribution. A) Proposed dual-inhibition model of gradient formation in differentiating hPSC colonies by Etoc *et al*^21^. B) Simplified mathematical representation of a generic dual inhibition model. C) Gradient formation of free BMP4 ligands as predicted by the model. D) Varying doses and colony sizes demonstrates the inability of the dual-inhibition model to generate a periodic Turing-like response in the free BMP4 distribution.

## Supplemental video legends

**Sup video 1**: **Reaction-Diffusion model of free BMP4 distribution in differentiating hPSC colonies**. The video shows the evolution of free BMP4 distribution in differentiating hPSC colonies in accordance with the BMP4-NOGGIN RD model in a differentiating colony of a 1000µm diameter.

**Sup video 2: Reaction-Diffusion model of predicts BMPi = 50ng/ml would not generate periodic patterns of free BMP4 distribution in differentiating hPSC colonies of 3mm in diameter**. The video shows the evolution of free BMP4 distribution in accordance with the BMP4-NOGGIN reaction diffusion model in differentiating hPSC colonies of 3mm in diameter when induced to differentiate with BMPi = 50ng/ml.

**Sup video 3: Reaction-Diffusion model of predicts induction of periodic waves of free BMP4 ligands distribution when hPSC colonies of 3mm in diameter are differentiated in BMPi = 200ng/ml**. The video shows the evolution of free BMP4 distribution in accordance with the BMP4-NOGGIN reaction diffusion model in differentiating hPSC colonies of 3mm in diameter when induced to differentiate with BMPi = 200ng/ml.

